# Structural Basis of Kinesin-1 Autoinhibition and Its Control of Microtubule-Based Motility

**DOI:** 10.1101/2025.07.15.665000

**Authors:** Md Ashaduzzaman, Yuqi Tang, Kyoko Okada, Stephen D. Fried, Richard J. Mckenney, Jawdat Al-Bassam

## Abstract

Kinesin-1 was the first identified microtubule-based motor protein that drives anterograde intracellular transport of diverse cargoes in eukaryotic cells. Improper regulation and kinesin-1 defects are implicated in multiple neurological disorders as well as pathogens hijacks kinesin-1 to deliver their cargoes. Despite its importance, the molecular mechanisms governing kinesin-1 regulation and activation remains poorly understood. Here, we report the cryo-EM structure of the autoinhibited kinesin-1 heterotetramer and validate it using crosslinking mass spectrometry. The structure reveals a 36-nm particle in which the kinesin heavy chains (KHCs) adopt a head to tail configuration, stabilized by asymmetrically arranged kinesin light chain (KLC) tetratricopeptide repeat (TPR) domains that bind across folded KHC coiled-coils and wedge in between the KHC motor domains. This architecture inhibits kinesin motility by constraining the dimeric motor domains in a configuration that is incompatible with processive movement. In addition, the structure shows that the KLC C-terminal helices occlude the TPR cargo binding interfaces, revealing a second layer of autoinhibition that directly blocks cargo engagement. Functional studies and structural modeling suggest that binding of regulatory factors, such as MAP7D3, compete with intramolecular KHC coiled-coil interactions, resulting in the unfurling of the autoinhibited structure and activating motor motility. These findings provide a molecular framework for understanding kinesin-1 regulation and its implications for intracellular transport.

## Introduction

Kinesin-1 (hereafter “kinesin”) is the founding member of a superfamily of motor proteins essential for intracellular transport along microtubule (MTs) (*1, 2*). Since its discovery in axonal transport studies, forty five genes encoding kinesin-related proteins have been identified in humans(*2*). As the most ubiquitous MT-based anterograde motor in eukaryotes, kinesin transports a diverse range of cargos, including mRNA, and organelles such as mitochondria and lysosomes, and nuclei, toward MT plus-ends(*3–6*). While its processive, hand-over-hand motility mechanism is well understood from studies of truncated constructs(*7, 8*), how the full-length motor is precisely regulated remains a central question.

In cells, kinesin exists predominantly as a heterotetramer composed of two kinesin heavy chains (KHCs) from one of three human isoforms (KIF5B, KIF5C, KIF5A) and two kinesin light chains (KLCs) from one of four human isoforms of KLC1-4(*9*). Each KHC is composed of a highly conserved N-terminal motor domain, followed by segments of conserved coiled-coils (CC0,CC1^KHC^,CC2^KHC^,CC3 and CC4) and terminating in C-terminal tail domains of distinct lengths(*10–12*). Each KLC is composed of N-terminal coiled-coils (CC1^KLC^,CC2^KLC^) connected to cargo binding <u>t</u>etratrico<u>p</u>eptide <u>r</u>epeat (TPR) domains via a conserved helical linker region (*13, 14*). The C-termini of KLCs contain variable sequence extensions that are subject to extensive alternative splicing(*15–17*). This structural complexity provides multiple points for cellular regulation, from autoinhibition via the KHC tail to cargo-dependent activation through the KLCs(*18*),(*19*).

The kinesin heterotetramer complex intrinsically folds into a compact, autoinhibited conformation (**Fig. 1A**), a state first inferred from hydrodynamic studies showing the native particle is roughly half the length of the motile motor(*14, 20, 21*). More recently, low-resolution structures from negative stain electron microscopy and cross-linking mass spectrometry (XLMS) visualized this “lambda” shaped particle (*22–24*). Nevertheless, the precise placement of individual KHC and KLC domains within this assembly has remains unclear.

**Fig. 1:**
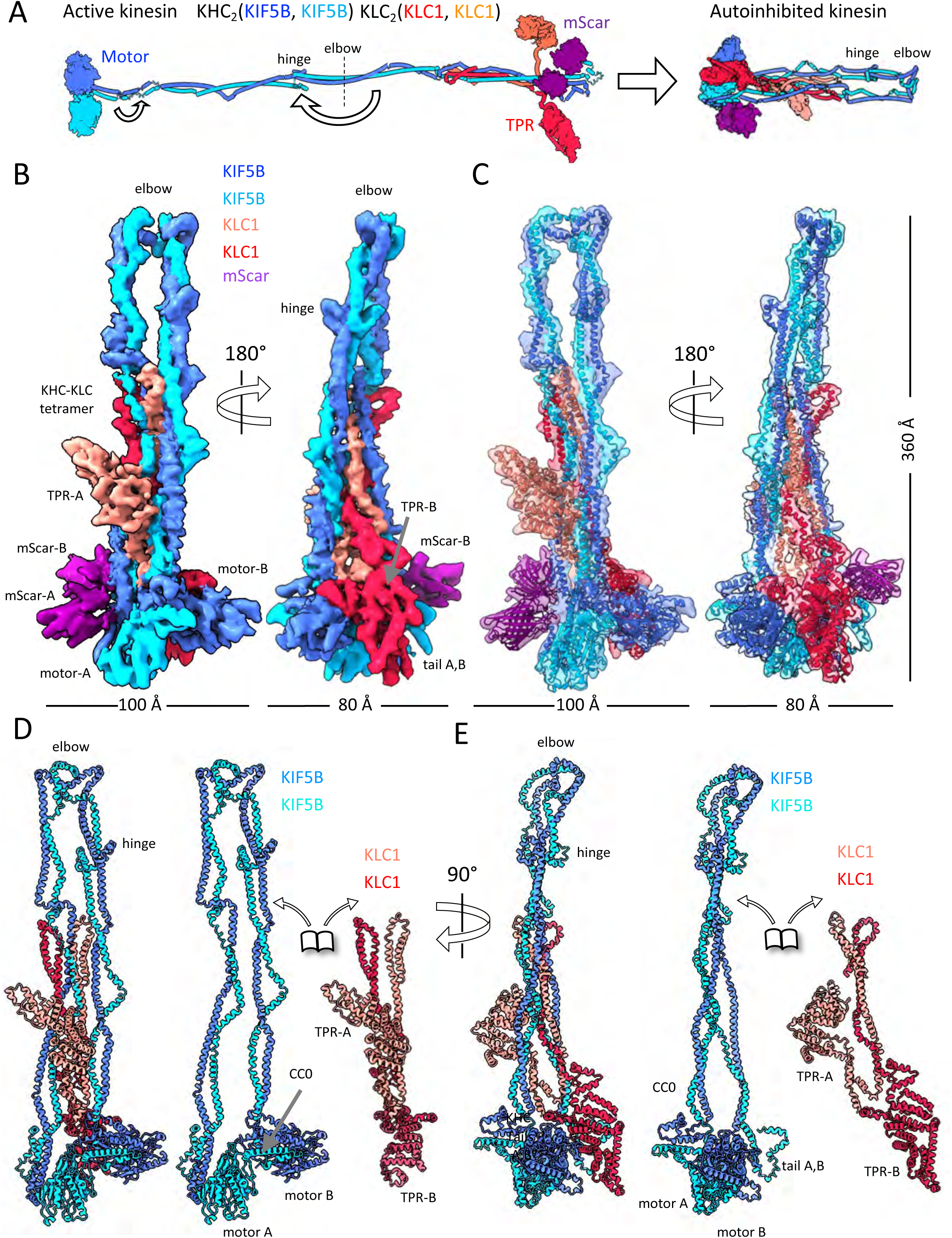
Cryo-EM structure of the autoinhibited kinesin KIF5B-KLC1 reveals the organization of the hetero-tetramer and the asymmetric subunit interactions in the lambda particle. A) A schematic showing the linear organization of kinesin heterotetramer with its KHCs (blue and cyan) and KLCs (red and orange). Arrows refer to the conformational folding events leading to the autoinhibited state structure described in this work. B) Two 180° rotated views of a segmented kinesin KIF5B-mScarlet-KLC1 heterotetrametric Cryo-EM map. The KHC subunits, KHC-A and -B, are colored blue and cyan, while the KLC subunits, KLC-A and -B, are colored red and salmon. Folded domain densities for the mScarlet (mScar-A, mScar-B) proteins fused on the KHC C-termini are colored purple, KHC motor domains (motor-A, motor-B) and KLC TPR domains (TPR-A, TPR-B) are also marked. C) Two 90° rotated views of the modeled (shown in ribbon) and segmented transparent kinesin KIF5B-mScarlet KLC1 heterotetramer cryo-EM map (transparent), colored as shown in A. D) Left, a ribbon model for KIF5B-KLC1 heterotetramer showing the asymmetric KLC-TPR interactions with KHC subunits without the mScarlet; Right, a dissociated view showing the isolated KLC (red and orange) and KHC (blue and cyan) dimeric subunits matching the orientation with left full heterotetramer model. E) 90° rotated view of ribbon models shown in D.

Beyond suppressing motility, the autoinhibited kinesin state must also control cargo engagement. Kinesin binds a vast array of cargoes through two main hubs: the KLC-TPRs recruit many reported kinesin cargoes of short conserved peptides found within cargo-adapter proteins that binding the concave interior interface of the crescent-shaped TPR domain(*25, 26*), while the KHC C-terminal tail regions bind cargo through adaptors such as JIP1, JIP3, tropomyosin and TRAK proteins(*27–29*). These adaptors regulate interactions with protein networks including Rab GTPases and kinases which promote signaling mediated activation of kinesin transport(*5, 30*). For proper regulation, these cargo-binding sites must be inaccessible in the inhibited state to prevent premature cargo loading and motor activation. Yet, it has remained unclear how the KLC TPR and KHC C-terminal domains are configured within the autoinhibited complex and how their functions are regulated.

Decades of research have provided key but sometimes paradoxical clues about the kinesin activation mechanism. The KHC C-terminal tail, containing a conserved isoleucine-alanine-lysine (IAK) motif, is a potent inhibitor of motor activity(*18, 23, 31*). However, while tail truncation produces a constitutively active motor, mutation of the IAK motif alone appears insufficient to fully activate processivity in the full-length complex(*22, 32*). A crystal structure of the kinesin motor domains with an isolated IAK-containing peptide reveals a 1:2 tail-to-motor complex, in which the peptide binds between the dimeric motor domains(*33*). However, studies that mutated the IAK motif in the full-length heterotetramer show that while the motif is required to repress MT binding, its removal is insufficient to strongly activate processive motility(*22, 32*). This suggests other factors are required, a role now largely attributed to cargo adapter proteins that bind to the KLCs(*26, 32*), and Microtubule associated protein 7 (MAP7) family proteins(*32, 34, 35*). MAP7 family proteins have also emerged as key regulators of kinesin activity both *in vitro* and in cells(*32, 34–37*). MAP7 proteins dramatically enhance kinesin’s MT association and are required for its activity in cells(*34, 35, 38*). This has led to a model of sequential activation, where MT-bound MAP7 first recruits and activates the motor, which in turn unmasks the KLCs for cargo binding(*38*). While this model explains many observations, the structural basis for this sequence of events—how the complex physically unfurls to release both motor and cargo inhibition—has remained a mystery.

To help resolve this, we determined an 8.6-Å cryo-EM structure of the autoinhibited human kinesin-1 heterotetramer. Our structure reveals a striking asymmetry in how the KLCs organize the complex, in contrast to the symmetrically intramolecularly folded KHCs. One KLC TPR domain acts as a clamp, stabilizing the folded KHC coiled coils, while the other acts as a wedge, locking the KHC tail and motor domains in a non-motile state. Furthermore, we uncover a novel mechanism of cargo-binding autoinhibition, in which C-terminal KLC helices fold back to sterically block the cargo-binding groove of the TPR domains. Together, these findings provide a comprehensive structural blueprint for how kinesin-1 co-regulates its motor and cargo-binding functions, establishing a physical framework for its sequential activation.

## Results

### Cryo-EM structure of kinesin in the autoinhibited state

We purified recombinant human kinesin heterotetramer, KIF5B-mScarlet and KLC1, as previously described(*32*), and optimized crosslinking conditions to enrich for the autoinhibited state (**Fig. S1A-B**). Mass photometry of sucrose density gradient purified fractions enabled the separation of the single heterotetramer (KHC_2_KLC_2_) population from the multimerized (KHC_4_KLC_4_) pools of kinesin heterotetramers (**Fig. S1C-F**). Extensive grid screening and vitrification yielded clean distributions of 40-nm single closed kinesin particles (**Fig. S2**). Full-size particles were only visible in relatively thick ice conditions, limiting the attainable resolution of our final cryo-EM reconstructions (**Fig. S2**). Initial 2D-classification revealed predominantly closed kinesin particles, with very few particles in the open conformation. 2D-classification also revealed flexibility in several regions of the closed 40-nm kinesin particle, particularly in its thinner coiled-coil regions (**Fig. S2**). Iterative 3D-classification yielded curated particle dataset and a 12-Å consensus kinesin map in which the folded coiled-coils, KLC-TPRs, mScarlet fluorescent proteins domains fused onto the KIF5B tails, and KHC-motor domains can be confidently identified by shape (**Fig. S3, top**).

To improve map quality and resolution across different regions, we sub-divided the kinesin particle into four overlapping regions and performed masked classification and refinement, yielding regional resolutions ranging from 5.6 to 8.6 Å (see Methods; **Fig. S3**). Nearly 80% of the kinesin particles showed flexibility in a density attributed to one KHC motor domain that interfaces between the KLC TPR domain and KHC-tail domains (**movie S1**). We refined two regions that include each of two KHC motor domains independently, while bound to neighboring elements to improve their local density (**Fig. S3; movie S1**). The refined subregional maps contained extensively overlapping features and comparable map qualities, allowing the clear alignment all subregional maps to obtain a consensus map at 5.6-8.6 Å resolution (**Fig. S3, Fig. S4A-F; movie S1**). The merged map of the full kinesin particle revealed clear helical secondary structure elements throughout, allowing model building of the kinesin assembled KHC and KLC coiled-coils, and folded domains in their heterotetrametric autoinhibited organization using either experimentally determined structures or Alphafold3 models (**Fig. 1B-C; Fig. S4; Fig. S5**). The model includes all KHC and KLC regions, except for the KLC N-terminal ∼20 residues, which were not resolved.

### Overall organization of autoinhibited kinesin structure

The cryo-EM structure reveals a 36-nm comet-shaped “lambda”(*23*) particle in which the symmetrical KHC dimer folds back on itself at the “elbow”, creating a turnaround zone for the second half of the KHCs **(Fig. 1B-C; movie S2)**. The two KLCs bind the KHC via their N-terminal coiled-coil regions but adopt asymmetrically arranged TPR domains: one TPR (termed TPR-A) stabilizes the folded coiled-coil interface, while the other (termed TPR-B) binds in between the motor domains and tails (**Fig. 1D-E**). The KHC coiled-coil containing regions fold back at the elbow junction to form the lambda particle(*23*), bringing the N-terminal KHC motor domains into proximity with the C-terminal KHC tails (**Fig. 1D-E**). Both KHC tails are both ordered, while fused, via short linker sequences, to two ordered mScarlet fluorescent β-barrel fold domains (termed mScar-A and mScar-B) (**Fig. 1B-C**).

The KHC and KLC helical, coiled-coils, globular motor and TPR domains generally matched their experimentally determined or AlphaFold predicted structures(*22, 23, 26*) (**Fig. S7**), but the structure revealed previously unknown intra- and inter-molecular interactions that assemble the autoinhibited state, which are described below in detail. The dimeric KHC consists of N-terminal motor domains (motor-A and motor-B), followed by a series of dimeric coiled-coil containing segments: CC0, CC1^KHC^, CC2^KHC^, CC3, and CC4 (**Fig. 1D-E; movie S2; Fig. 2A-B**). The dimeric KLCs consist of two N-terminal KHC-binding KLC coiled-coil regions (CC1^KLC^ and CC2^KLC^) followed by linker-helical regions (L-H: residues 164-198), and a cargo binding TPR domain (residues 199-479). The structure reveals KLC regions following the TPR domain; KLC C-terminal helical regions include—Helix-1 (H1: residue 480-509), Helix-2 (H2: residues 510-522), self-cargo (SC: residues 522-535), and Helix-3 (H3: residues:535-575). These KLC regions are all well-ordered and are crucial for maintaining the kinesin autoinhibited state (see below; **Fig. 2; movie S3)**.

**Fig. 2:**
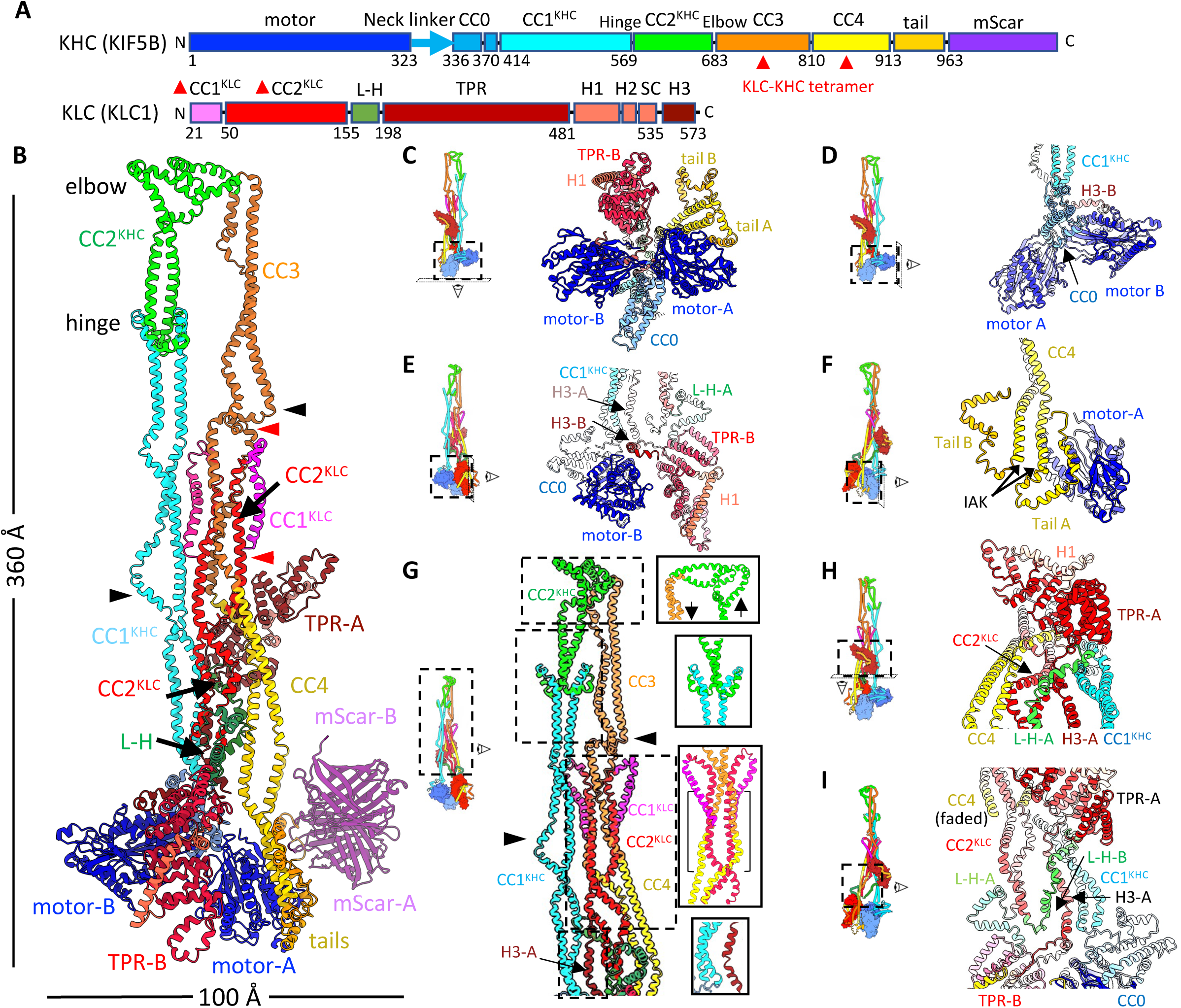
The organization of the KHC and KLC domains in the kinesin autoinhibited structure. A) Domain maps of human KIF5B and KLC1. The KHC (KIF5B) domains include the motor domain, neck linker, CC0, CC1^KHC^, CC2^KHC^, CC3, CC4 and tails. The mScarlet (mScar) fused onto the C-terminus is shown. The KLC (KLC1) includes: CC1^KLC^, CC2^KLC^, L-H, TPR domain, and the H1, H2, Self-cargo (SC), H3 helical regions. Boundaries are indicated in residue numbers, below, and each are colored to match colors for their models shown in B and C. B) The kinesin-1 KIF5B-KLC1 heterotetramer model with domains colored as shown in panel A. KLC-TPR-A and -B are shown in red and dark red, respectively. Black Arrowheads refer to CC1^KHC^ and CC3 regions with super-twisted or untwisted coiled-coil regions, respectively. C) Bottom view showing the KHC motor-A, -B, tail-A, B and the KLC-TPR-B D) Side view showing the KHC dimeric motor-A, B domains and the packed CC0 and CC1^KHC^ domains E) Side view showing the KHC motor-A interacting with KLC-TPR-B and the self-folded state of CC1^KHC^ and CC4. F) Side view showing the KHC motor-B interacting asymmetrically with tail-A and tail-B which emerge from CC4. The IAK motifs, labeled with arrows, lie at the end of CC4. G) Side view showing the coiled-coil regions of kinesin including CC1^KHC^, CC2^KHC^, CC3, CC4, CC1^KLC^, CC2^KLC^ and H3. The KHC-CC N- and C-termini are labeled. H) Top-tilted view showing the KLC-TPR-A bound to the two KHC halves stabilizing CC1^KHC^ folding onto CC4 I) Side view of the structure, with CC4 faded, showing the CC1^KLC^ and two directions of the L-H-A, -B extending toward the KLC-TPR-A and TPR-B

### A structural model for autoinhibition of motor activity via extensive intramolecular self-folding

The overall 36-nm structure is composed of the KHC dimer folding back on itself, aided by the KLC domain interactions (**Fig. 1D-E**). This lambda-shaped fold of the KHC comprises a “forward” arm (CC0, CC1^KHC^, CC2^KHC^) and a “return” arm (CC3, CC4), connected by a novel U-shaped structure, the “elbow”(*23*), which facilitates the 180° turnaround at the C-terminus of CC2^KHC^ (**Fig. 2B-I**). This anti-parallel arrangement holds the N-terminal motor domains in an orientation incompatible with MT binding, providing a direct structural explanation for autoinhibition, showing extensive restrictions on KHC motor motile free interactions. The entire KHC assembly is further stabilized by the asymmetrically folded KLC dimer where TPR-A binds against the antiparallel KHC coiled-coil regions, while TPR-B bifurcates the motor domains in the head-to-tail KHC interaction interfaces (**Fig. 1C**; **Fig. 2B-C**).

Several interactions lock the KHC motor domains into this inhibited conformation. The neck coiled-coil (CC0) forms a dimeric four-helix bundle that is packed between the two motor domains, orienting them perpendicular to the particle’s long axis (**Fig. 2C-E**). This state is reinforced by the KHC tails (tail-A and tail-B) (**Fig. 2F**), which emerge from the CC4 return arm to interact with only one motor domain (motor-A) (**Fig. 2F**). This asymmetric 2:1 tail-to-motor interaction is consistent with biochemical findings⁵¹, but is structurally distinct from contacts observed in prior models developed from isolated KHC fragments or KHC motor-tail fusion MT-bound complexes(*33, 39*). Finally, the previously designated ‘hinge’ region(*40, 41*) at the junction of CC1^KHC^ and CC2^KHC^ (**Fig. 2G**) is clearly observed and is in good agreement with recent computational models and negative stain EM data(*22, 23*), validating another key feature of the KHC coiled-coil fold. Despite the lower resolution, the cryo-EM maps suggests parts of CC1^KHC^ and CC4 coiled-coils adopt super-twisted or under-twisted coiled-coil conformations, respectively, (black arrowheads, **Fig. 2B,G)** suggesting the potential local relief of the molecular stress built-up in KHC coiled-coil-containing regions upon self-folding; this is consistent with previous findings that kinesin-1 KHC changes coiled-coil propensity(*42*). Similar changes in coiled coil twist are observed in the dynein stalk domain and the BICD2 dynein activator(*43, 44*).

The autoinhibited kinesin structure revealed a surprisingly asymmetric organization of the dimeric KLC subunits (**Fig. 2E, H, I**). CC1^KLC^ folds around each KHC CC3-CC4 in anti-parallel manner leading to CC2^KLC^, which runs parallel to CC3-CC4 (**Fig. 2G**). The N-termini of the CC2^KLC^ helices form a tetrameric parallel coiled-coil bundle with KHC CC3-CC4, resulting in heterotetramer assembly (Red Arrowheads, **Fig. 2A-B, G**). This produces a parallel eight helical bundle forms in this region sandwiched between CC1^KHC^-and CC4 (**Fig. 2B, G**). Following this tetrameric KHC-KLC assembly zone, the remaining CC2^KLC^ 90-residues assemble into homo-dimeric coiled-coils that terminate with two linker-helical regions (L-H-A, L-H-B; **Fig. 2E, G-I**). Although, L-H -A and L-H -B regions emerge from the symmetric CC2^KLC^ dimer, they fold in opposite directions toward the asymmetrically positioned KLC-TPRs (**Fig. 2B, E, H-I)**. KLC-TPR-A bridges alongside the two KHC-KLC coiled-coil arms (**Fig. 1**; **Fig. 2H)**, while KLC TPR-B wedges between the two motor domains and closely interacts with motor-B (**Fig. 1**; **Fig. 2B**). The total buried surface area by KHC-KLC interaction is approximately 5500 Å and represent mostly coiled-coiled and TPR interaction interfaces (**Fig. S6**).

The autoinhibited kinesin cryo-EM structure is a significant departures from previous models generated by AlphaFold or from negative-stain EM data(*22*) (**Fig. S7**). Key differences include the overall organization of the KHC coiled-coils, the binding interfaces of the KLC TPR domains, and the orientation of the motor domains. Notably, while the general position of the elbow is consistent with that reported by Tan et al.(*22*), its structure is fundamentally different. Instead of the sharp turn predicted by AlphaFold, our structure reveals a broad, U-shaped bend spanning ∼40 residues (KIF5B: residues 640-686) that guides the KHC back upon itself **(Fig. 2B, G)**. Furthermore, our model provides the first complete structural view of the autoinhibited states of the neck coiled-coil (CC0) (**Fig. 2C-D)** and the C-terminal tails (**Fig. 2F)**, clarifying their crucial roles in kinesin autoinhibition and regulating activation.

### Crosslinking mass spectrometry validates the interfaces in kinesin autoinhibited structure

To understand the multi-subunit interactions within the autoinhibited kinesin in relation to the cryo-EM structure, we carried out crosslinking mass-spectrometry (XLMS) of same autoinhibited kinesin heterotetramer used in our cryo-EM structures, but using BS3 crosslinker (**see Methods**). In total, we identified 592 unique crosslinks in the kinesin sample, including 279 intra-KIF5B crosslinks, 106 intra-KLC1 crosslinks, 115 inter-subunit crosslinks between KIF5B and KLC1, and 92 crosslinks involving mScarlet (**Fig. 3C**; **Fig. S8**). Of these, 572 crosslinks could be mapped onto the kinesin autoinhibited cryo-EM model, with 297 (52%) satisfying the 35 Å distance cut-off (**Fig. 3A, C**). These XLMS data are consistent with the kinesin autoinhibited structure model, both in overall architecture (**Fig. 3C**) and in terms of specific inter-domain contacts (**Fig. 3D-I**). Since our cryo-EM structure determination suggested flexibility in various regions of the kinesin autoinhibited structure (**movie S1**), we also evaluated crosslinks with extended distances (35-50Å) and mapped those onto the structure in comparison to the shorter distance crosslinks (**Fig. S9**). With these additional crosslinks, 68% of the crosslinks are consistently mapped onto the structure (**Fig. S9C**).

**Fig. 3:**
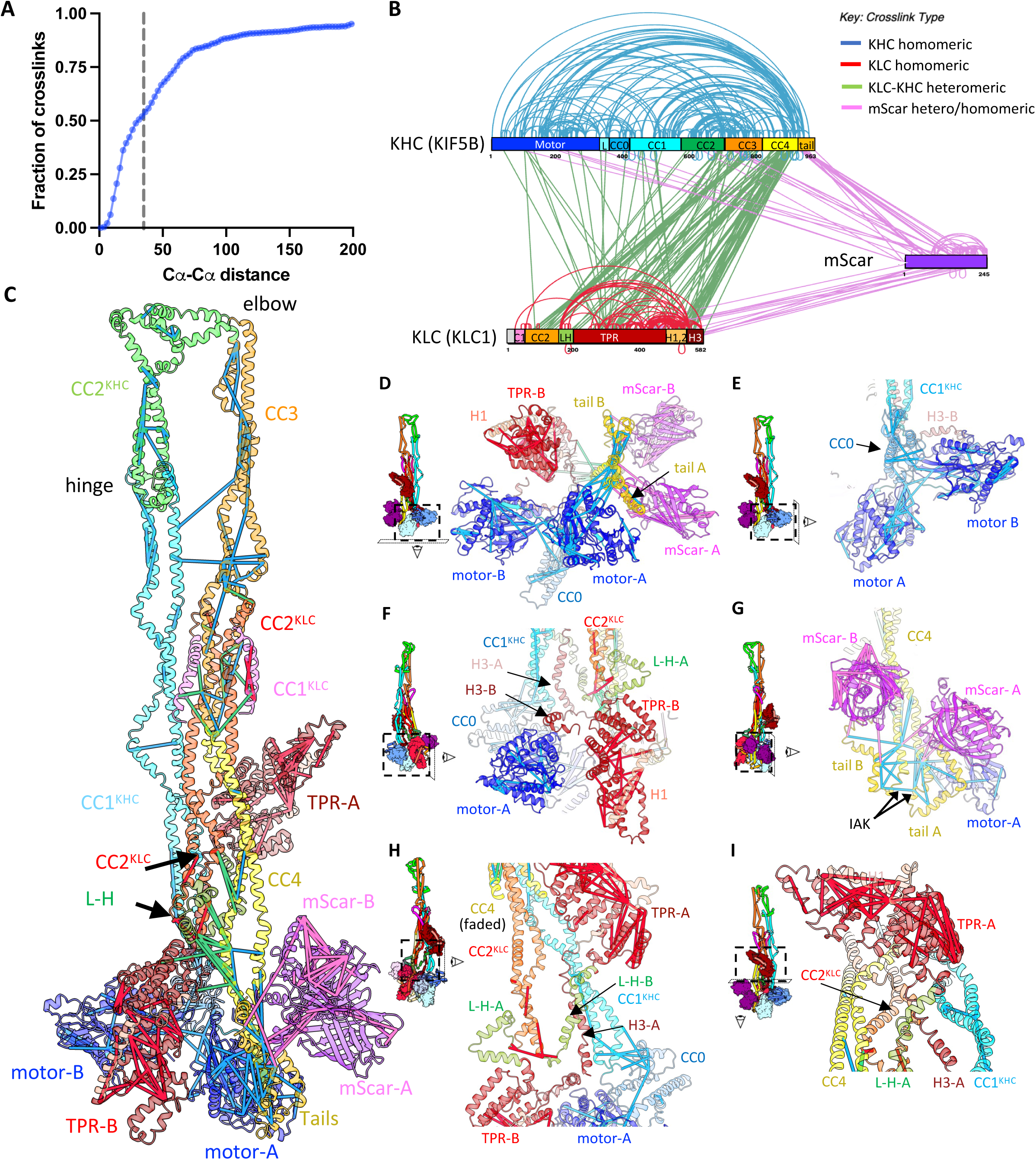
Crosslinking mass spectrometry validates the kinesin autoinhibited structure. A) Cumulative distribution function (CDF) plot showing the fraction of the 572 crosslinks whose Cα-Cα distance is less than the indicated distance. The dotted line denotes the 35 Å cutoff. B) A linear domain map of KIF5B (KIF5B; top left) with the fused mScarlet (mScar; center right) and KLC1 (KLC1; bottom left) with internal crosslinks highlighted in blue, pink and red, respectively. The heteromeric crosslinks between KIF5B and KLC1 are shown in green and the heteromeric crosslinks between mScarlet and KIF5B and KLC1 are shown in pink. C) Crosslinks identified in kinesin-1, illustrated in the context of the cryo-EM structural model of kinesin KIF5B-mScarlet-KLC1 heterotetramer. In total, 572 crosslinks were successfully mapped onto the kinesin cryo-EM structure, of which 297 with Cα-Cα distances less than 35 Å are displayed. The view matches that in Fig. 2B. F D) Bottom view showing the KHC motor-A, -B, tail-A, B and the KLC-TPR-B. The view matches that in Fig. 2C. E) Side view showing the KHC dimeric motor-A, B domains and the packed CC0 and CC1^KHC^ domains. The view matches that in Fig. 2D. F) Side view showing the KHC motor-B interacting with KLC-TPR-B and the self-folded state of CC1^KHC^ and CC4. The view matches that in Fig. 2E. G) Side view showing the KHC motor-A interacting asymmetrically with tail-A and tail-B which emerge from CC4. The view matches that in Fig. 2F. H) Top-tilted view showing the KLC-TPR-A bound to the two KHC halves stabilizing CC1^KHC^ folding onto CC4. The view matches that in Fig. 2H. I) Side view of the structure, with CC4 faded, showing the CC1^KLC^ and two directions of the L-H-A, -B extending toward the KLC-TPR-A and TPR-B. The figure matches that in Fig. 2I.

Close-up views of the crosslinks, reveal numerous short-range crosslinks within both folded KHC and KLC domains, showing a strong consistency with the domain contacts resolved in our cryo-EM structure (**Fig. 3C-I**). For example, 48 crosslinks were identified within the KHC motor domains; 2 and 6 crosslinks were observed connecting the KHC motor domain to CC0 and CC4-tail, respectively (**Fig. 3C-E**). In addition, we noticed two short-range (<20 Å) crosslinks that connecting CC1^KLC^ and CC3 domains (**Fig. 3C**). All these observations match the anti-parallel fold-back of the KHC coiled-coils. 14 crosslinks were observed between the KHC CC3-CC4 and CC2^KLC^, which confirms the dimerization of the KHCs and KLCs and resulted tetramer bundle (**Fig. 3C**). We also identified three short-range crosslinks connecting the KLC L-H and TPR region to the KHC CC4 region (**Fig. 3C**). Together with the observation of numerous long-distance crosslinks between the KLC TPR region and nearly all regions of KHC, these findings are consistent with our asymmetric model for the two TPR regions within the kinesin heterotetramer (**Fig. 3C**). We observe extensive crosslinks within each mScarlet domain, and between the mScarlet domains and KHC motor, CC0, CC4 and tail domains (**Fig. 3C, D, G**). Extended distance crosslinks (35-50Å) mostly map to interfaces between the KLC TPR-B and KHC tail domains, and map to the interface between KLC TPR-A and the CC1^KHC^ and CC4 and CC2^KLC^ (**Fig. S9D-F**). This suggest that the KHC tail and coiled-coil interfaces with the KLC TPR are probably flexible, consistent with flexibility analyses (**movie S1**).

We also compared our XLMS data to previously published KIF5B-KLC1 kinesin heterotetramer XLMS dataset by Tan et al^12^ (**Fig. S8 E**). Mapping their identified crosslinks onto our cryo-EM model revealed that 147 of 233 crosslinks (63%) fall within the 35 Å distance cut-off, indicating a high level of consistency with the model (**Fig. S8B, E**). Notably, their dataset did not include the KIF5B-fused mScarlet. Our XLMS data provide confidence in the assigned location of mScarlet by showing five short-range crosslinks between the mScarlet and KHC CC4-tail domains (**Fig. 3C, D, G**). In addition, we detected numerous long-range crosslinks linking mScarlet to other domains, including the KHC motor, coiled-coil, and tail regions, as well as the KLC TPR and helical CC2^KLC^ and L-H regions (**Fig. 3C; Fig. S9**). Consistent with this model XLMS crosslinks show the Lys922 in IAK motif crosslinks to residues in KHC motor (residues 166,213) as well as crosslink to residues in the KHC tail (residues 903,908,933) which lie in other helices of the W-hairpin tail (**Fig. 3G**). Hence these XLMS data serve as orthogonal validation of the cryo-EM structural model, including the location of the KIF5B fused mScarlets and the location of the KHC tails that were previously improperly placed in the negative stain EM or AlphaFold models(*22*) (**Fig. S7**).

### Dimeric KHC C-terminal tails bind to one motor domain in the autoinhibited state

The KHC C-terminal tails, which are crucial for forming the autoinhibited state(*45*),(*22, 23*), are revealed to be ordered in the structure as W-shaped helical hairpin domains (**Fig. 2B, C, F**). Each tails forms a helical W-hairpin emerging as dimers at the end of CC4. Similar, W-shaped hairpin motifs are found on many other proteins with two-strand antiparallel coiled coil and are crucial for binding other proteins or RNA(*46*). The dimeric KHC tails interact with the motor domains in a strikingly asymmetric fashion. Only one tail (tail-A) directly engages a single motor domain (motor-A), binding near the α2 (P-loop) and α3 (Switch II) helices. Our model suggests that the conserved Ile-Ala-Lys (IAK) motifs are positioned at the junction of CC4 and the first tail helix (**Fig. 2 C, F**). The KHC tails also make contacts with the KLC-TPR-B on the other side, as evidenced by XLMS crosslinks (**Fig. S9**). This new KHC tail-motor binding site is consistent with differential ADP to ATP nucleotide exchange between the two motor domains(*47*). The IAK motif junction regulates dimeric KHC tail orientation to motor-A. The second tail (tail-B) remains unbound from motor-A, but may shield an auxiliary MT interaction site at the end of CC4, which is consistent with finding that KHC tails lower the KHC CC4 affinity for MTs (*48*). This asymmetry in KHC tail to motor interactions are further evidenced by the staggered positions of the C-terminally fused mScar-A, mScar-B; mScar-A fused to the bound tail-A is positioned ∼10 Å lower than the mScar-B fused to the unbound tail-B, suggesting that binding to motor-A constrains and tightens the tail-A structure (**Fig. 1-2**).

This model of the KHC tail-motor interaction differs significantly from previous structures derived from co-crystals of isolated motor domains and tail peptides or from cryo-EM maps of MT-bound motor-tail fusion constructs(*33, 39*). Our autoinhibited kinesin cryo-EM structure demonstrates that the native inhibitory KHC tail-motor domain contacts is critically dependent on the structural context—specifically, anti-parallel orientation of the dimeric KHC CC4-tails to one motor domain (motor-A) and to a KLC-TPR (TPR-B) (**Fig. 2B-C**). Structural comparisons of our model to the crystal structure of the isolated motor-tail show that if motor-A aligned with then motor-B is found in different conformations while only binding of tail-peptide between the two motor domains together (**Fig. S10A-B)**. While in those structures the KHC tail-peptide binds near ß-4 of both isolated tai-motor domain structures, this differs from our observation of one tail binding near α2 and α3 of motor-A in the autoinhibited kinesin structure (**Fig. S10B**). The new tail binding site revealed by our cryo-EM structure lies on the opposite side of the motor, next to its P-loop (**Fig. S10C**). The KLC TPR and KHC-CC0 interactions with motor-A and motor-B occlude the space for any of the states in which motor B binds the KHC tail as observed in the isolated crystal structure or MT decorated cryo-EM maps of those constructs (**Fig. 2G**). Furthermore, the KHC C-terminal tail to motor domain interactions are likely preserved even in the absence of KLCs, as suggested by previous negative-stain EM studies on KHC dimers²⁴, highlighting the tail to motor interaction to as a fundamental feature of KHC autoinhibition.

### Asymmetric binding of the KLC TPR domains to either to KHC coiled-coiled or a KHC motor domain

Despite recent insights into autoinhibited kinesin(*22, 23*), the precise positioning of the KLC TPR domains has remained an open question. Our cryo-EM structure resolves this by revealing that the two KLC-TPR domains bind in a surprisingly asymmetric manner to the symmetric KHC dimer (**Fig. 1D-E**; **Fig. 2B, C, E, H**; **Fig. 4A**). Instead of engaging equivalent sites, each TPR domain bind to a distinct region of the KHC, adopting unique roles. One TPR domain (TPR-A) bridges across the CC1^KHC^, CC3-CC4 and CC2^KLC^ anti-parallel coiled-coil bundle (**Fig. 2H**), while the other (TPR-B) separates the KHC motor-A-B domains, possibly constraining their movement (**Fig. 2B, C, E)**.

**Fig. 4:**
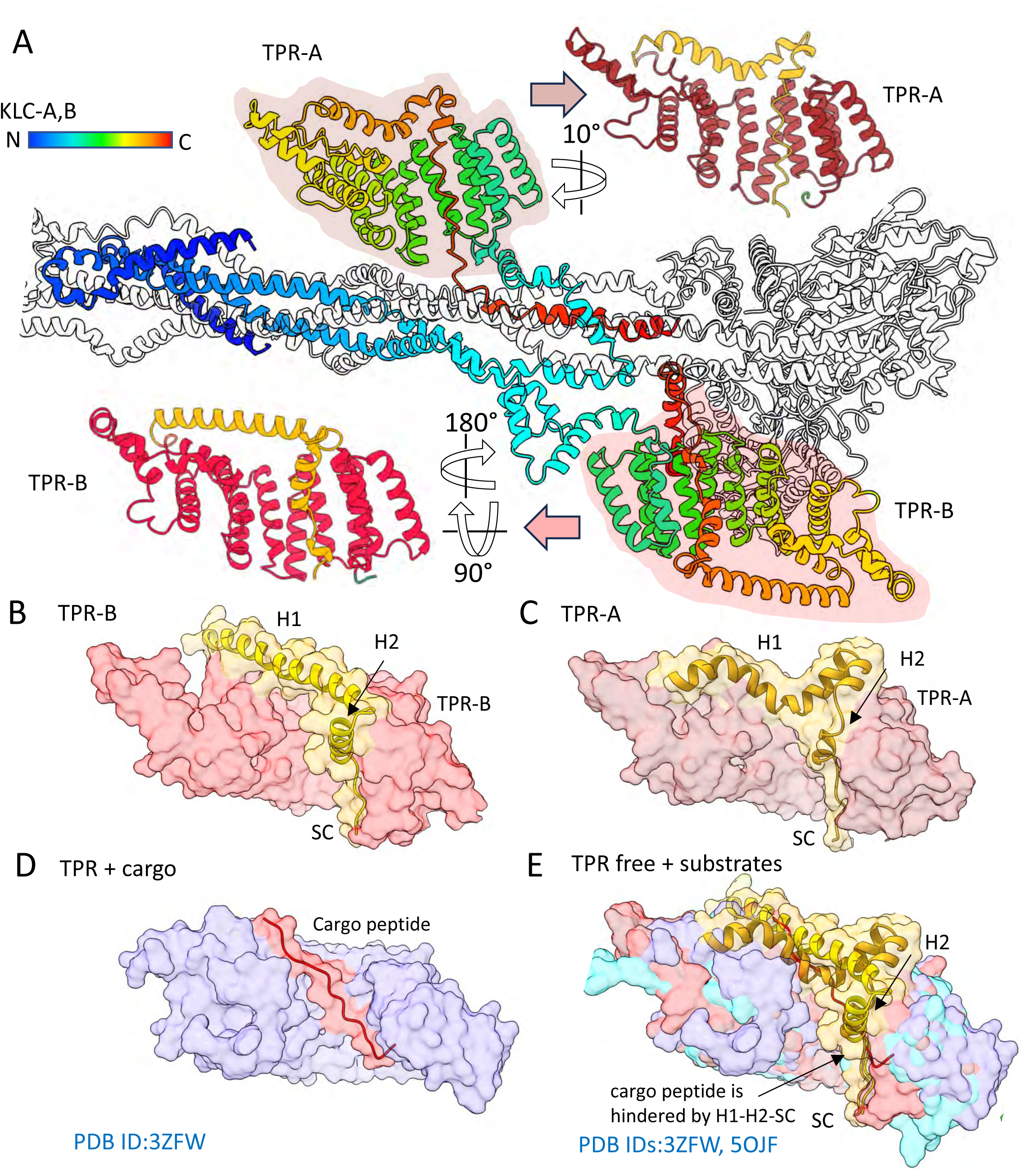
The topology and self-regulation of the KLC subunits in lambda kinesin autoinhibited structure. A) A general view of the kinesin structure with KHC colorless and KLC is shown in rainbow gradient from the N- to C-terminus. Each TPR domain is shown and its orientation transformed via arrows to orient them in a comparable manner. The isolated KLC-TPR models show the role of the KLC C-terminal helical regions, colored in gold, autoinhibiting access to the cargo binding concave surface of the TPR domains. B) A view of the KLC-TPR-B domain with the helical regions of the TPR fold shown in surface representation with the KLC-C-terminal helical regions shown in ribbon form. C) A view of the KLC-TPR-A domain in the same orientation described in B D) A view of the KLC-TPR structure (PDB-ID:32FW) with the Y-acidic motif cargo peptide bound to the concave surface of the KLC-TPR(*26*). E) An overlay of the KLC-TPR structure (brown and salmon) in two states described here and the free (cyan; PDB ID:5OJF) and Y-acidic motif cargo-peptide bound form (blue; PDB-ID:3ZFW) overlaid. The SC and H2 in autoinhibit the TPR domain and sterically hinder cargo peptide binding.

TPR-A stabilizes the folded KHC architecture by bridging the CC1^KHC^-CC2^KHC^ and CC3-CC4 arms (**Fig. 2H)**. It uses its narrow surface, composed of inter-helical repeat turns, to bind the KHC coiled-coils, maintaining them together in their folded conformation (**Fig. 1**; **Fig. 2**). This interface is constrained by the TPR-A N and C-termini bound to KHC coiled-coils (see below; **Fig. 2B, I**). This interaction is critical for maintaining the overall integrity of the ‘lambda’ particle. This region is a major source of flexibility around these coiled-coils (**movie S1**). In contrast, TPR-B is positioned as a triangular wedge between motor-A and motor-B domains, on the opposite side of the dimerized KHC neck or CC0. The positioning of TPR-B between the motor domains may explain the elevated autoinhibition of kinesin KHC in the presence of KLC(*32, 49*) and is consistent with the report of motor domain bifurcation by TPR(*21*). TPR-B interacts closely with motor-B, using the helices of its TPR repeats 2-5 to lock motor-B in a restricted state (**Fig. 2C, E**). The amino acid sequences at these unique KLC-KHC interaction interfaces are highly conserved across species (**Fig. S11**), underscoring their functional importance.

Beyond constraining the KHC head-to-tail fold in the autoinhibited state, our cryo-EM structure suggests another mechanism for KLC autoinhibition (**Fig. 4**). In both KLC subunits, ordered C-terminal helical regions—H1, H2, and a self-cargo (SC) motif—fold back to occupy the concave, cargo-binding face of their respective TPR domains, sterically preventing adapter protein engagement (**Fig. 4B-E**). This complex, asymmetric arrangement is accommodated by distinct conformations of the KLC N- and C-terminal linker helices (L-H and H3). For example, the linker following TPR-A (H3-A) binds along CC1^KHC^, while the linker following TPR-B (H3-B) binds across the top of motor-B (**Fig. 2E, I)**. Our work provides the first complete view of all KLC helical segments within the context of the fully autoinhibited kinesin structure.

### The KLC TPRs are autoinhibited by C-terminal helical sequences

Previous studies proposed that KLC is autoinhibited via an N-terminal ‘LFP’ sequence that competes with cargo proteins for binding to the TPR domain(*50, 51*) (**Fig. 4D**). We find that the concave cargo-binding groove of each TPR domain is occupied by a series of helices (H1, H2, and SC motif) that are C-terminal to the TPR domain itself (**Fig. 2**; **Fig. 4A-C**). H1 acts as a connector that folds back from the TPR domain, positioning H2 and the SC motif to sterically block the interface for Y- or W-acidic cargo motifs (**Fig. 4B-D**). This self-inhibited state is further locked in place by the most C-terminal helix, H3, which anchors the KLC to adjacent KHC domains (CC1^KHC^ or motor-B). A subtle difference seen between the two KLC TPRs is H1-A is that more bent toward the concave face of TPR-A, while H1-B is extended straight along TPR-B (**Fig. 4B-C**). Our structure confirms that the KLC TPRs exist in an autoinhibited state but reveals a fundamentally different mechanism for cargo inhibition, driven by the KLC C-terminal helical elements (**Fig. 4E**).

Our findings directly contrast with a previously proposed N-terminal ‘LFP’ mechanism(*50*). We attribute this discrepancy to the different experimental contexts: our structure represents the full-length heterotetramer, whereas earlier studies utilized isolated KLC proteins with truncated C-termini that lacked the inhibitory helices we observe here (*50*). The context of the full complex is therefore essential for revealing the mechanism of KLC autoinhibition.

### Restrictions of the kinesin motor domains in the autoinhibited structure

While it is known that full-length kinesin motility is strongly repressed(*32, 40*), the precise structural mechanism has been unclear. Our structure reveals that the dimeric KHC motor domains are locked into a tightly constrained, non-processive configuration through a multi-point locking mechanism (**Fig. 5A-B**). This is achieved by specific interactions with one KHC C-terminal tail (tail-A), a KLC TPR domain (TPR-B), and the neck coiled-coil (CC0). tail-A binds directly to motor-A, while TPR-B separates the two motor domains, binds KHC tails, and tightly engages motor-B. Simultaneously, the CC0 dimer is packed into a unique conformation against the two motor domains at their dimerization junction, and further restricts their flexibility from opposite side to the TPR-B and KHC-tails (**Fig. 2B-F**; **Fig. 5A-B**).

**Fig. 5:**
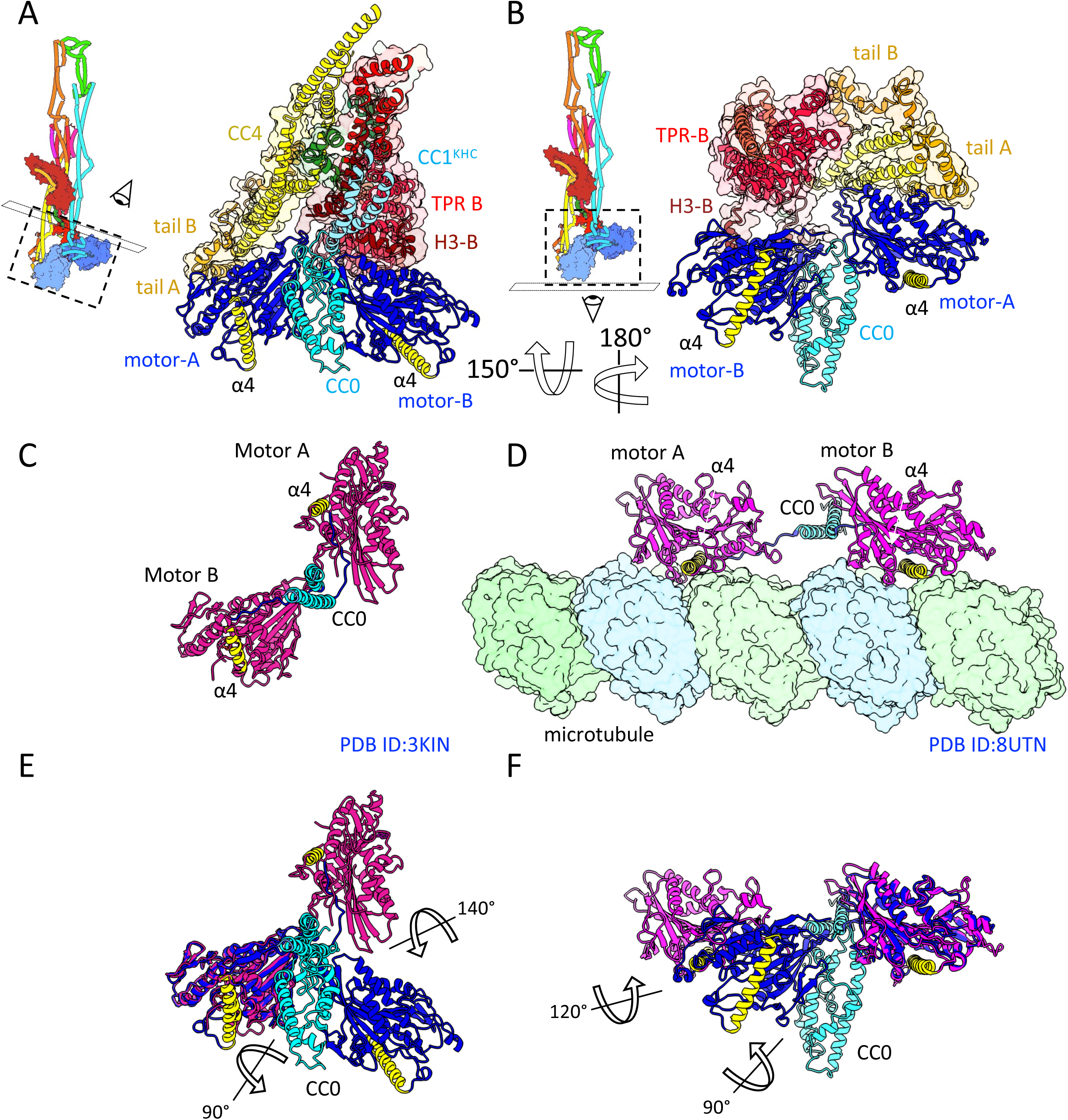
The kinesin motor domains are restricted from motility by the lambda autoinhibited kinesin structure interactions. A, B) two views of the Kinesin structure showing the KHC motor A, B (blue) bound by the KHC tail-A, -B (orange and gold) and the KLC-TPR domain (red) and CC0 (cyan) is packed beneath CC1^KHC^. The MT binding site of each motor domain is highlighted with α4 helix shown in yellow. C) A view matching the orientation shown in panel A of the dimeric kinesin motor domain crystal structure(*52*) (PDB ID: 3KIN) showing the motor domains (magenta) and the CC0 (cyan). The MT binding site of the motor domain MT is highlighted with α4 helix shown in yellow. D) A view matching the orientation shown in panel A of the dimeric KIF1A motor domain cryo-EM model along the MT protofilament(*53*) (PDB ID:8UTN) showing the motor domains (magenta) and the CC0 (cyan). The MT binding site of the motor domain is engaged with the MT, and this binding site is highlighted with α4 helix shown in yellow. E) Overlay of view of the CC0, motor-A and -B domains shown in A with those shown in C. F) Overlay of view of the CC0, motor-A and -B domains shown in B with those shown in D.

To understand the functional consequences of this locked arrangement, we compared it to motile or unbound kinesin structures, including the MT-bound KIF1A dimer and the free rat kinesin motor dimer crystal structure(*52, 53*). This comparison reveals a dramatic reorientation: one of the motor domains in our autoinhibited structure is twisted by 120°-140° around its neck linker junction relative to its position in the motile states (**Fig. 5C-F**). As a direct result, the MT-binding interfaces (the α4 helices) of the two motor domains are severely misaligned and cannot engage the MT lattice sites simultaneously (**Fig. 5E-F**). Therefore, our structure demonstrates that autoinhibition physically prevents processive motility by locking the motor domains in a conformation where the hand-over-hand stepping mechanism is impossible. Also, our flexibility analyses (**movie S1**) and classifications (**Fig. S2; Fig. S3**) show that motor-B which does not interact with tail-A or B is more flexible which likely suggests it maybe be the motor domain able to initiate MT binding in the autoinhibited state.

### Map7D3^CT^ binding to CC1^KHC^ destabilizes the kinesin autoinhibited state

MAP7 family proteins are critical for kinesin function *in vivo*(*34, 36*). While it is established that MAP7 enhances kinesin’s MT association, its direct role in activating motility has been ambiguous(*34–36*). Some studies suggest full activation requires a cargo-adapter protein(*32*), while others observed increased motility that could be attributed to a higher MT landing rate rather than true activation(*34, 35*). More recently, the paralog MAP7D3, which binds kinesin more tightly, was shown to enhance motility even in truncated K560 constructs (KIF5B, residues 1-560), comprised only of the motor-domains-CC0 and CC1^KHC^, suggesting a more direct activation mechanism^43^.

To investigate how MAP7D3 activates kinesin, we first used AlphaFold3 to model the interaction between the MAP7D3 C-terminal domain (MAP7D3^CT^:residues 476-876) and the KHC(*41, 54, 55*). The model predicted that residues 680-729 of MAP7D3 form a four-helix bundle with the CC1^KHC^ region, an interaction that would sterically clash with the autoinhibited fold and disrupt critical stabilizing contacts (**Fig. S12, Fig. 6A**). To test this hypothesis, we performed single-molecule TIRF microscopy assays with kinesin heterotetramer (KIF5B-mScarlet KLC1) with and without MAP7D3 to assess whether MT binding is necessary for MAP7D3^CT^ interactions. As expected, the kinesin heterotetramer was strongly autoinhibited in the absence of MAP7D3^CT^(*32, 34–36*). In stark contrast, the addition of MAP7D3^CT^ triggered a ∼12-fold increase in the number of processive runs which showed a higher motility velocity than the paused kinesin without MAP7D3 (**Fig. 6B-C**). These results demonstrate that MAP7D3^CT^ can relieve kinesin autoinhibition, consistent with the idea that MAP7D3^CT^ binding CC1^KHC^ disassembles the autoinhibited state, providing direct functional validation for our structural model.

**Fig. 6:**
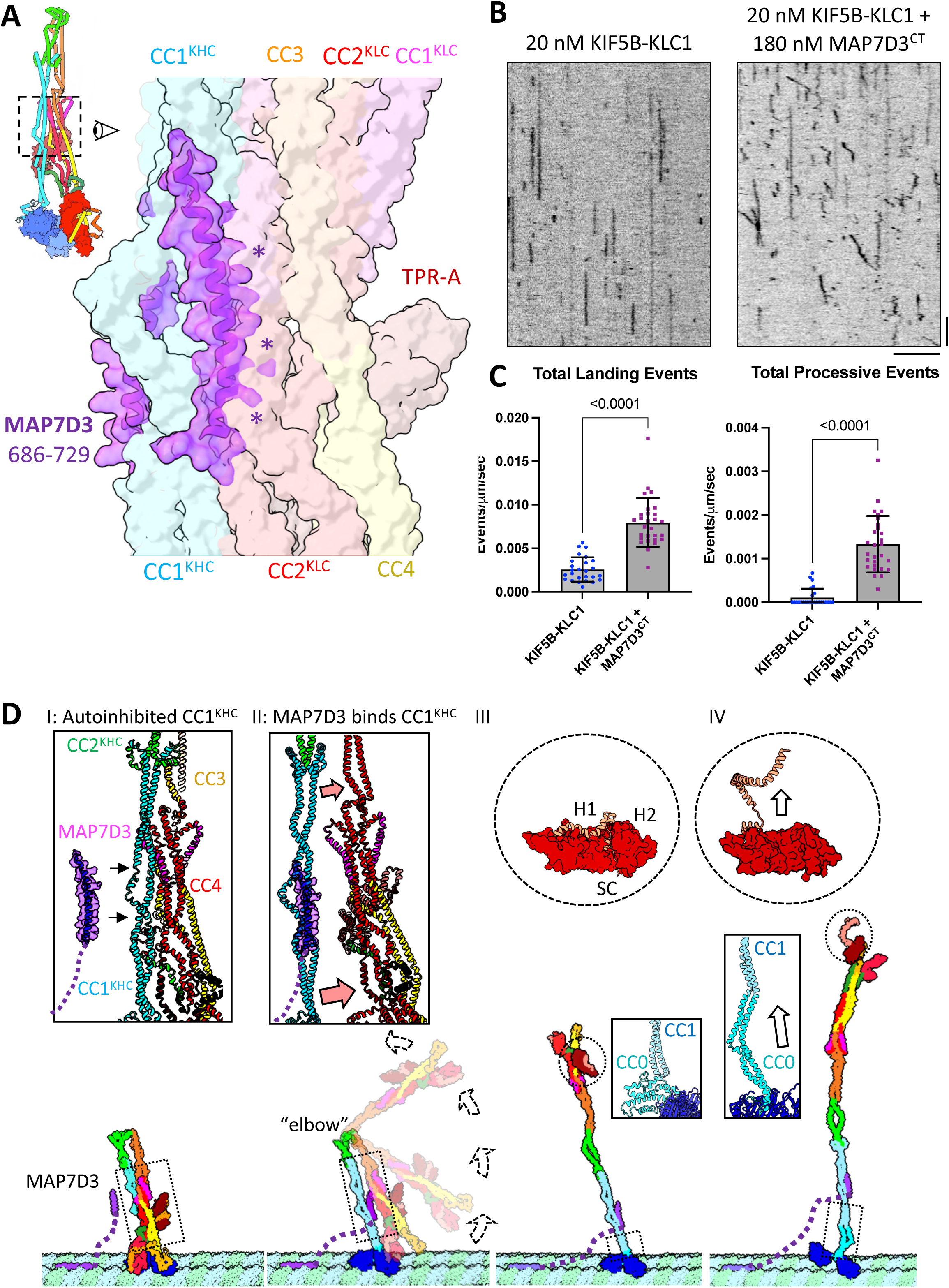
MAP7D3^CT^ is predicted to bind CC1^KHC^, sterically competing with the kinesin autoinhibition interfaces, increasing MT landing rate and enabling kinesin motility along MTs. A) An overlay of the Alphafold3 model of MAP7^CT^ (residues 686-728) (purple envelope and ribbon) binding to CC1^KHC^ onto the CC1^KLC^-CC2^KLC^-CC4 kinesin autoinhibited structure (faded surface colors) revealing MAP7 binding induces steric hindrance with the CC2^KLC^ and CC4 interface on CC1^KHC^. Details in (**Fig. S8; Fig. S12**). B) Kymographs of Kinesin reconstitution with MTs visualized by TIRF microscopy. Left panel autoinhibited kinesin binds along MTs (left) and leads to transient and static association events. Right panel, in the presence of MAP7D3^CT^, kinesin shows enhanced landing rates and activation of processive motility toward MT plus ends (left end). C) Left panel, Quantitation of MT landing events for kinesin in the absence and presence of MAP7D3^CT^. Right panel, Quantitation of processive motility events. Student’s t-test was used to compare datasets leading to P values provided. D) Model illustrating the MT-mediated transition and MAP7D3-driven activation of kinesin through disruption of the autoinhibited lambda conformation. Domains are color-coded as in Fig. 2. Panels I–IV (bottom) depict the overall kinesin conformation and its transitions on the MT lattice in response to MAP7D3 binding. Top panels provide close-up views of specific domains and regions undergoing structural changes during activation. Activation begins when MT-bound MAP7D3 **(I)** binds to the CC1^KHC^, causing a steric clash with KHC-CC3-CC4 and CC2^KLC^ that destabilizes and unfurls the autoinhibited complex **(II)**. This large-scale rearrangement sequentially **(III)** releases the motor domains for motility by unraveling the CC0 neck coiled-coil **(III-IV, central panels)** and **(IV)** unmasks the KLC TPR cargo-binding sites by displacing the C-terminal inhibitory helices **(III-IV, top panels)**, rendering the motor fully active for transport.

This activation mechanism also explains how MAP7D3 enhances cargo binding, as shown in previous work(*38*). Our model suggests this occurs because disrupting the autoinhibited complex by interacting with CC1^KHC^ and thus liberates the KLC TPR domains from CC1^KHC^, which is consistent with recent finding from Shukla *et al*(*56*). The autoinhibited kinesin interacts with the MT and encounters MAP7D3^CT^ leading to the relief of autoinhibition to release kinesin to undergo processive runs. Therefore, MT bound MAP7D3 functions as a dual-action activator: it sterically disrupts the autoinhibitory interface to directly trigger motility, which in turn unmasks the KLCs to promote the recruitment of cargo adapters(*38*).

## Discussion

Kinesin-1 is the founding and most extensively studied member of the kinesin superfamily. For decades, research on kinesin-1 has defined its hand-over-hand motility mechanism(*3, 18, 57*), yet the molecular basis of its spatiotemporal regulation has remained poorly understood. It is well established that the kinesin heterotetramer is autoinhibited via intramolecular interactions(*40, 45, 48*), but a complete structural picture of this state has been missing and somewhat contradictory(*21, 32, 40, 41, 58*). Deletion of the KHC C-terminal tail region or newly identified KHC elbow regions leads to enhanced kinesin motility(*23, 40*). Recent work has converged on a model of sequential activation, in which factors like MAP7 first activate motility, which in turn permits the loading of cargo via the KLC TPR domains(*32, 34–36, 38*). However, the structural basis for this coordinated process—how the motor is inhibited from both moving and binding cargo, and how activators could reverse this—has been a central and unresolved question.

Our cryo-EM structure and XLMS analysis of the fully autoinhibited kinesin provides a structural basis of its autoregulation. We show that the complex is a precisely organized assembly in which long-range interactions to create a dual-locked state. First, motor activity is suppressed by a network of asymmetric contacts in the autoinhibited state: the two KLC TPRs either bind the together anti-parallel folded CC1^KHC^ and CC4/CC2^KLC^ and separate the dimeric KHC motor-A and motor-B domains (**Fig. 1**; **Fig. 2; movie S2; movie S3**). Flexibility analyses (**movie S1**) and the XLMS studies (**Fig. 3**; **Fig. S9**) support these TPR interactions and suggest they have dynamic states. In conjunction with the KHC C-terminal tails, the KLC TPRs extensively restrict the free motility of the KHC motor domains which is essential for processive hand over hand motility (**Fig. 1**; **Fig. 2**; **Fig. 5**). Second, and unexpectedly, cargo binding by the KLC TPR is independently blocked by the C-terminal KLC helical segments, in which H1, H2 and SC occupy the concave TPR cargo-binding groove, while H3 binds distant elements that are in close proximity to maintain the autoinhibited state (**Fig. 2**; **Fig. 4**). This mechanism of cargo peptide occlusion is fundamentally different from previously proposed models (**Fig. 4**)(*26, 51*). This dual-inhibited kinesin architecture provides a comprehensive blueprint fail-safe for how kinesin maintains robustly an ‘off’ state, while also revealing distinct structural points of regulation that can be targeted by activators to sequentially unlock its motility and cargo loading. It also provides a map to interpret the functional variation of the paralogues of KHC and KLC with distinct sequences at their C-termini.

### A structure-based model for kinesin activation and the sequential loading of cargo

Our structural and biochemical data converge on a comprehensive model for the sequential activation of kinesin, integrating MAP7-mediated recruitment with the relief of the autoinhibition of both motility and cargo-binding **(Fig. 6)**. We propose a structural transition cascade begins when MT-bound MAP7D3^CT^ recruits an autoinhibited kinesin heterotetramer, increasing its local concentration on the MT **(Fig. 6D, I)**. Consistent with previous reports(*34, 35*) and our AlphaFold predictions (**Fig. S12; Fig. 6A**), the MAP7D3^CT^ likely binds to CC1^KHC^. The binding of MAP7D3^CT^ higher affinity of the MAP7D3^CT^ to CC1^KHC^ sterically clashes with the CC3-CC4 and CC2^KLC^, destabilizing the interfaces between the two folded halves of the KHC/KLC coiled-coil interfaces (**Fig. 6D, top insert I-II**). This initial disruption triggers a large-scale conformational rearrangement, leading to the unfurling of the full kinesin heterotetramer, nearly doubling its length from ∼36 nm to ∼72 nm, which releases inhibitory constraints on the motor domains—including the unravelling of the CC0 neck coiled-coil **(Fig. 6D, III-IV, middle panels)**—to permit hand-over-hand motility. Concurrently, this global unfolding liberates the KLC TPR domains via the release of KLC-H3, causing the KLC C-terminal inhibitory helices H1, H2 and SC to dissociate from the TPR cargo-binding groove. This final step unmasks the cargo-binding sites, rendering the now-motile kinesin fully competent for cargo-adapter engagement **(Fig. 6D, III-IV, top panel)**. The packed-up state of the KHC neck (CC0) coiled-coil likely also unravels due to the absence of the autoinhibition interactions with the KLC TPR and KHC tail domains, leading to its release and the freeing of the KHC motor domains to undergo hand over hand motility (**Fig. 6D, III-IV middle panels**).

In conclusion, elucidating the intramolecular interactions that govern kinesin-1 autoinhibition has been a long-standing challenge in the cytoskeletal motor field. Our work provides a structural answer, revealing a comprehensive blueprint for how motor activity and cargo engagement are co-regulated. By demonstrating how a network of asymmetric, long-range interactions creates a dual-locked state, this model establishes a clear foundation for future mutational studies to dissect these mechanisms both *in vitro* and *in vivo*. Furthermore, it provides a powerful framework for exploring functional differences in autoregulation across the entire kinesin superfamily.

## Supporting information

no links necessary

## Author Contribution statement

MA carried out biochemical studies, prepared cryo-EM grids, collected cryo-EM data, determined and refined all structures, built and refined all models, co-wrote and co-revised manuscript. YT carried out XLMS studies, prepared figures, co-revised manuscript. KO carried out biochemical preparations and co-revised manuscript. SDF interpreted XLMS data, co-revised manuscript. RJM carried out biochemical studies and co-wrote and co-revised manuscript. JAB conceived the project, provided grant support, supervised and advised all co-authors, carried out biochemical experiments, prepared figures, wrote and revised manuscript.

## Data Availability statement

The kinesin cryo-EM regional maps, assembled full map, and model coordinates were submitted to the EMDB and PDB with the following accession numbers EMDB: EMD-71730, EMD-71731, EMD-71732, EMD-71733, EMD-71734, and PDB ID: 9PMB. The mass spectrometry proteomics data have been deposited to the ProteomeXchange Consortium via the PRIDE partner repository with the dataset identifier PXD073177 and 10.6019/PXD073177 and provided the crosslinks in the supplementary file Data S1.

## Competing interest statement

The authors declare no competing interests

## Acknowledgement

We thank Dr Michael Cianfrocco (University of Michigan, Ann Arbor) for early support, advice in early stages and encouragement throughout the project. We thank Dr. Kyoko Chiba (Tohoku University) for the generation of kinesin reagents, advice and experimental support. We thank Dr. Bharti Singal (Stanford University Cryo-EM Center) for preliminary studies. We thank Dr Camille Scott and the UC-Davis High performance computing core facility (HPCCF) for computational support. We thank Dr Gant GW Luxton (Molecular Cellular Biology, University of California, Davis) for comments on the manuscript. We would like to thank the Dr Ed Eng, Dr Eugene Chua at the NIH National Center for Cryo-electron microscopy Access and Training (NCCAT) for providing advice and help with sample preparation and for initial data collection. Large scale Cryo-EM data were collected at NCCAT and the Simons Electron Microscopy Center located at the New York Structural Biology Center, supported by the NIH Common Fund Transformative High Resolution Cryo-Electron Microscopy program (U24 GM129539, and NIGMS R24 GM154192) and by grants from the Simons Foundation (SF349247) and NY State Assembly. JAB and RJM are supported by the NIH (R01GM110283 and 1R35GM158334 to JAB and R35GM124889 to RJM). SDF acknowledges support from the Sloan Foundation and YT is supported by the NIH (R01GM129301).

## Materials and Methods

### Protein expression and purification kinesin-1 heterotetramer and MAP7D3^CT^

For Kinesin-1 expression, KIF5B-mScarlet and KLC1 were cloned into pACEBac1/pIDS vector, DH10MultiBac (Geneva Biotech) were transformed as recombined plasmid to generate bacmid as previously described(*32*). Baculovirus was prepared by bacmid transfection using Cellfectin II reagent (Thermo Fisher Scientific) followed by two cycles of amplification leading to P2 virus. For protein expression, 400 mL of Sf9 cells (2-3 × 10^6^ cells/mL) were infected with 4 mL of P2 virus and cultured for 65 h at 27C. Cells were harvested and resuspended in 25 mL of lysis buffer (50 mM HEPES-KOH, pH 7.5, 150 mM KCH_3_COO, 2 mM MgSO_4_, 1 mM EGTA, 10% glycerol) along with 1 mM DTT, 1 mM PMSF, 0.1 mM ATP and 0.5% Triton X-100. After incubating on ice for 10 min, the lysates were centrifuged at 15,000 X g for 20 min at 4C. The resulting supernatant were subject to affinity chromatography in which the supernatants were pumped over a column of Streptactin XT resin (IBA) for 1 h at 4C. The columns were then washed with excess amount of lysis buffer to remove unbound material, and the proteins were eluted in lysis buffer containing 100 mM D-biotin. Eluted proteins were further purified using Size exclusion chromatography using TSKgel- a Phenomenex BioSep 5 mm SEC-s4000 500 Å Column with a size of 600 x 7.8 mm as previously described(*32*).

For MAP7D3^CT^ expression, MAP7D3^CT^ was cloned into bacterial expression vector and transformed into BL21-CodonPlus (DE3)–RIPL E. Coli (Agilent, Santa Clara, CA). Cultures of BL21 expressing MAP7D3^CT^ were grown at 37°C in Luria broth with kanamycin (50 μg/ml) until an optical density at 600 nm of 0.6. Protein expression was induced with 0.2 mM IPTG at 18 °C overnight. Bacterial cultures were centrifuged at 5000 x g and pellets were frozen. Bacterial pellets were thawed on ice and resuspended in purification buffer (PB:50 mM tris-HCl (pH 8.0), 150 mM KCH3COO, 2 mM MgSO4, 1 mM EGTA, and 5% glycerol) freshly supplemented with 1 mM PMSF, 0.1 mM ATP, NucA nuclease, and protease inhibitor mix (Promega, Madison, WI). For purifying Map7D3ct, 1 mM dithiothreitol (DTT) was also added during this step. Bacteria were lysed by passage through an Emulsiflex C3 high-pressure homogenizer (Avestin, Ottawa, ON, Canada) and the subsequent by addition of 1% Triton X-100 for 5 min on ice. Lysed cells were then centrifuged at 22,769g for 20 min at 4°C. or affinity purification, the clarified lysates were incubated with resin as follows. Map7D3ct was incubated with Streptactin XT resin (IBA Lifesciences, Göttingen, Germany) for 1 hour at 4°C and washed with PB. Map7D3^CT^ was further purified using a HiTrap Capto S cation ex-change chromatography column (Cytiva) equilibrated in HB buffer [35 mM Pipes- KOH (pH 7.2), 1 mM MgSO4, 0.2 mM EGTA, and 0.1 mM EDTA, pH 7.1]. Bound proteins were eluted with a 45- ml linear gradient of 0 to 1 M KCl in the HB buffer.

### Cryo-EM sample preparation and data collection

Kif5B-mScarlet-KLC1 at 1 mg/mL was analyzed on sucrose density gradient (**Fig. S1A**). We then tested crosslinking with 0.01-0.08% (v/v) glutaraldehyde at room temperature for 30 minutes and analyzed on SDS-PAGE gel to find optimal crosslinking concentration (**Fig. S1B**). The sample was prepared for cryo-EM sample with 0.06% (v/v) glutaraldehyde with GraFix(*59*) system using 20 mM HEPES pH 7.2, 100 mM KCl, 1 mM MgCl_2_, 5 mM B-ME, 1 mM EGTA and loaded onto 10-40% (v/v) sucrose gradient with 1 mg/mL Kif5B/KLC1 and ran for 16 hours at 4 C at 40000 rpm (**Fig. S1A**). Sample was manually collected with a pump at 10s interval of ∼200 μL. Fractions containing Kif5B-mScarlet KLC1 was then neutralized with 1 mM Tris-HCl pH 7. Crosslinked samples were dialyzed to remove sucrose with 20KDa MWCO Amicon filters and concentrated with 100 KDa MWCO centrifugal filter. GraFix crosslinked samples were also tested on Superose 6 5/150 gel filtration column (**Fig. S1C**) and ran on SDS-PAGE showing the fully crosslinked Kinesin (**Fig. S1D**). The crosslinked samples were analyzed for their average masses using a Refeyn Mass Photometer (Refeyn Ltd.) revealing a mass consistent with pure heterotetramers (**Fig. S1E**) and was further used to curate sample showing oligomers (**Fig. S1F**).

Quantifoil R1.2/1.3, 300 mesh Au grids were plasma cleaned with H2/O2, 50 W, for 30 sec and Vitrobot blotting conditions were set at 4°C and 100% humidity. Kif5B-mScarlet-KLC1 sample with detergent FOM (0.12x-1x CMC) (VitroEase™, Thermo-Fisher) at 3 mg/mL was pipetted to the grids and blotted 3-3.5s and plunge-freeze in liquid ethane.

For Cryo-EM datasets collection, G2 Krios microscope (ThermoFisher) at 300 kV equipped with Falcon IV detector (ThermoFisher) at 130,000x magnification was set and data acquisition was done using Leginon(*60*) beam-image shift(*61*) with 7s exposure with total dose of ∼48 e-/Å at pixel size 0.926 Å with defocus −0.8 µm to −2.5 µm at 130,000x magnification and filtered through a Selectris 20 eV energy filter. Total four datasets of 55674 movies were saved as Electron-event-representation (EER)(*62*) files at image dimensions of 4096×4096 pixel.

### Cryo-EM structure determination

Each EER-Movie dataset was imported into RELION(*63*) separately and Motion-corrected with EER fractionation of 32 using RELION’s own implementation with B-factor of 150 and patches of 5×5 with no binning. CTF estimation was done using CTFFIND-4.1(*64*). Initial picking was done using Laplacian -of-Gaussian (LoG) in RELION with 100×450 diameter and extracted in 200-pixel box at 3.704 Å/pixel followed by improved picking using negative stain map and Topaz. Multiple 2D-, 3D-classification and 3D-refinement steps helped to remove junk, broken or open kinesin particles leading to 36-nm comet-shaped ‘lambda’ particle structures from each of the datasets in which the particle has a wide globular organization at one end and an elongated narrow region at the other end. Iterative, 2D-classifications were done in cryoSPARC(*65*) and particles were re-extracted in RELION(*63*) and re-classified in cryoSPARC(*65*) and ∼4 millions particles were re-picked with Topaz(*66*) in RELION(*63*). To accurately center these elongated ‘lambda’ particles, we used an iterative strategy of picking particles followed by T=0 classification(*67*) (**Fig. S2**). Aligned particles were iteratively refined, classified, and polished (**Fig. S3**) to accumulate 664938 particles extracted to 1.852 Å/pixel. Global consensus 3D-refined structures were determined using iterative 3D-refinement and 3D-classification in large box formats. Density for the sub-regions of the consensus cryo-EM map were further improved by re-centering, re-extraction, and local refinement leading to 5.6-8.6 Å resolution (**Fig. S3; Fig. S4**). Re-centering and re-extraction into 150-pixel box with 1.852 Å/pixel followed by local refinement used for TPR-A and part of CC1^KHC^ and CC4 and KLC-CCs with 5959495 particles. 3D-classification without alignment shows single class with 103108 particles with both motor in closed conformation which were then 3DFlex(*68*) and local refined in cryoSPARC(*65*) to resolve the motor-tail with mScarlets-KLC TPR-B (**Fig. S3**). These refined subregional maps were combined by overlaying onto the consensus map, leading to full kinesin ‘lambda’’ particle density map to 5.6-8.6 Å resolution throughout the structure. Model placed map guided by local (CC1-CC4, motors, KLCs) and a 36 nm flex refined maps initially from 26k particles (CC2-CC3) were used to generate 15 Å lowpass filtered map in EMAN from SBGrid software packages(*69*) a and then was used to align the CC2-CC3 turnaround after 60 Å lowpass filtering before refinement to solve a 12 Å map which was then followed by 3D classification, refinement and 3D variability analysis separating 30746 particles which was local refined at 300 pixel box at 1.852 Å/pixel. Local resolutions were estimated using PHENIX(*70*) (**Fig. S3; movie S1**). Cryo-EM data collection, single particle data processing, and final model statistics are shown **Table I**.

**Table I:**
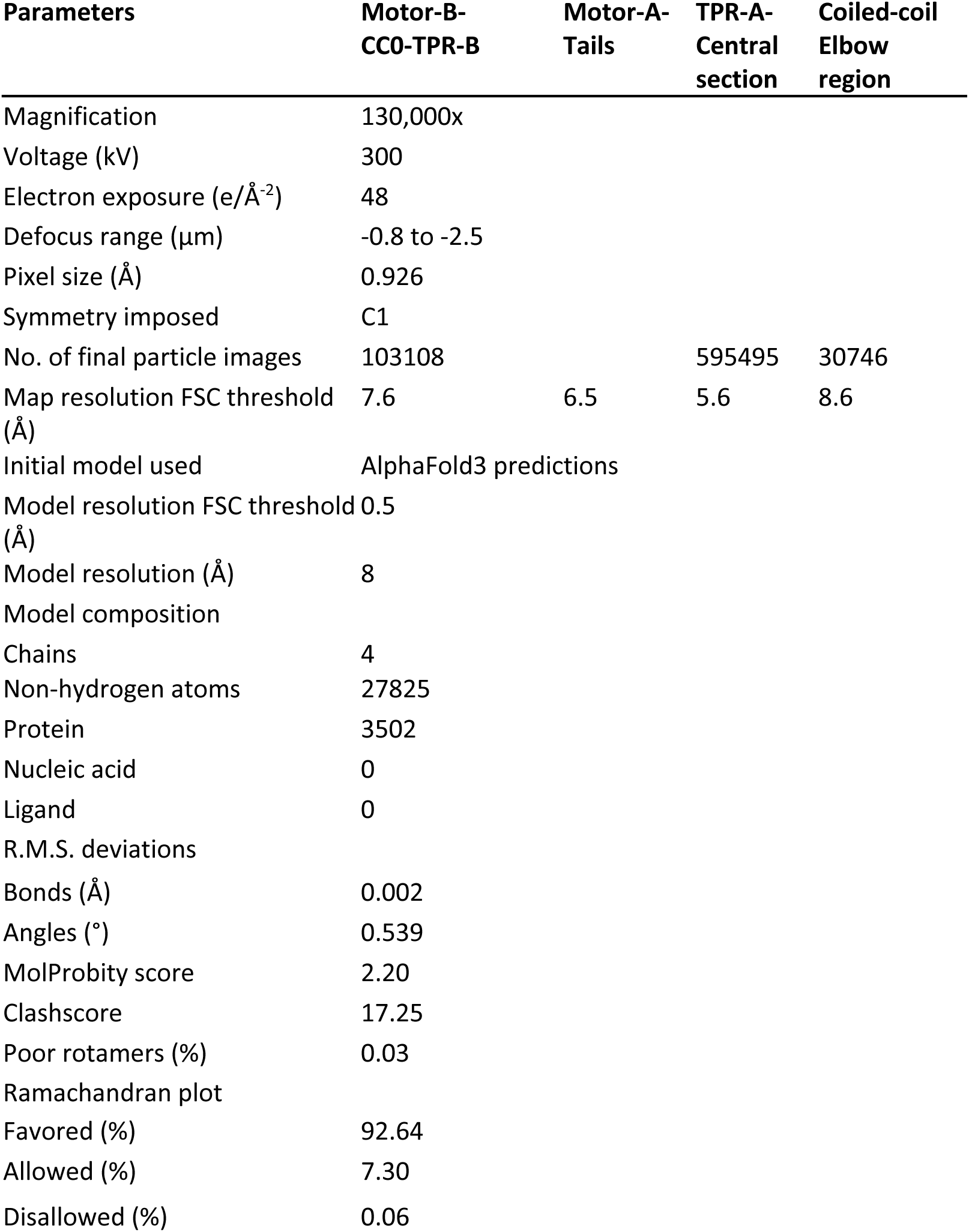
Cryo-EM data collection, structure determination and model building statistics.

| Parameters | Motor-B-CCO-TPR-B | Motor-A-Tails | TPR-A-Central<br>section | Coiled-coil<br>Elbow<br>region |
| --- | --- | --- | --- | --- |
| Magnification | 130,000x |  |  |  |
| Voltage (kV) | 300 |  |  |  |
| Electron exposure (e/Å<sup>-2</sup>) | 48 |  |  |  |
| Defocus range (μm) | -0.8 to -2.5 |  |  |  |
| Pixel size (Å) | 0.926 |  |  |  |
| Symmetry imposed | C1 |  |  |  |
| No. of final particle images | 103108 |  | 595495 | 30746 |
| Map resolution FSC threshold (Å) | 7.6 | 6.5 | 5.6 | 8.6 |
| Initial model used | AlphaFold3 predictions |  |  |  |
| Model resolution FSC threshold (Å) | 0.5 |  |  |  |
| Model resolution (Å) | 8 |  |  |  |
| Model composition |  |  |  |  |
| Chains | 4 |  |  |  |
| Non-hydrogen atoms | 27825 |  |  |  |
| Protein | 3502 |  |  |  |
| Nucleic acid | 0 |  |  |  |
| Ligand | 0 |  |  |  |
| R.M.S. deviations |  |  |  |  |
| Bonds (Å) | 0.002 |  |  |  |
| Angles (°) | 0.539 |  |  |  |
| MolProbity score | 2.20 |  |  |  |
| Clashscore | 17.25 |  |  |  |
| Poor rotamers (%) | 0.03 |  |  |  |
| Ramachandran plot |  |  |  |  |
| Favored (%) | 92.64 |  |  |  |
| Allowed (%) | 7.30 |  |  |  |
| Disallowed (%) | 0.06 |  |  |  |

### Crosslinking mass spectrometry (XL-MS) of kinesin assemblies

Kinesin KIF5B-mScarlet-KLC1 heterotetramer was crosslinked in nearly identical conditions to cryo-EM, as described above, with 0.5-2 mM BS3 at 4°C overnight using sucrose density gradient. The crosslinked samples were the denatured (8 M urea), reduced (5 mM DTT), alkylated (15 mM IAA), quenched with DTT, digested with LysC (1:100 w:w ratio), diluted 4-fold, digested with trypsin (1:50 w:w ratio), desalted (Sep-Pak C18), and vacuum-dried. The sample was analyzed using an UltiMate3000 UHPLC system coupled to a Q-Exactive HF-X Orbitrap. Peptides were resuspended and loaded onto PepMap 100 C18 column with Solvent A (0.1% formic acid in water) and solvent B (0.1% formic acid in 98% ACN). Then the peptides were separated using PepMap RSLC C18 column with a linear gradient from 5% to 25% solvent B over 100 min, then to 45% over 25 min, and to 90% over 5 min. For LC-MS/MS data acquisition, the Q-Exactive HF-X performed MS1 scans at 120,000 resolution (350-1500 m/z), AGC target of 3 × 10^6^, and 50 ms max IT. The top 10 precursors (z = 3-8) were isolated (1.4 m/z window) and fragmented using stepped NCE (32±3). MS2 scans were at 60,000 resolution (200-2000 m/z), AGC target of 1 × 10^5^, and 150 ms max IT. Dynamic exclusion was set to 45 s and in-source CID at 10 eV.

RAW files were converted and recalibrated using the xiSEARCH preprocessing pipeline python script. The recalibrated MGF file was searched with xiSEARCH 1.8.9 for crosslinked peptides. Search parameters were used as following: MS1 mass tolerance, 6 ppm; MS2 mass tolerance, 10 ppm; allowed maximum number of missed cleavages, 4; minimum peptide length, 6. Crosslinks were searched based on the modifications at Lys and N-term (preferred), modifications at Ser, Thr, and Tyr were also allowed with lower priority. Carbamidomethylations (+57.021464 Da) on Cys were enforced as fixed modifications, oxidations (+15.99491463 Da) on Met and phosphorylations on Ser/Thr/Tyr (Sp: + 166.9984 Da; Tp: +181.0140 Da; Yp: +243.0269 Da) were allowed as variable modifications. FASTA file containing KIF5B, KLC1 and mScarlet was applied in the search separately. Search results were filtered in xiFDR 2.3.10 at the residue pair level to an FDR of 5%. The boost function was enabled between residue pairs, and the rest of the settings were default. The mzid file generated from xiFDR was uploaded onto xiVIEW and PDB file generated from cryo-EM study was imported for crosslink visualization. **Fig. 3C-I** and **Fig. S8 B,D** were generated by exporting the filtered crosslinks from xiVIEW and rendered with the cryo-EM ribbon model in ChimeraX(*71*) with XMAS(*72*). The connectogram in **Fig. S8C** was generated with xiVIEW without filtration. The mass spectrometry proteomics data have been deposited to the ProteomeXchange Consortium via the PRIDE partner repository with the dataset identifier PXD073177 and 10.6019/PXD073177 and provide a table of all unique crosslinks in the supplementary file Data S1.

### Model building and XLMS data visualization

Local refined subregional maps were aligned and a ChimeraX(*71*) vop map was created for full-length Kinesin-1 heterotetramer (**Fig. S4A-D**). AlphaFold3 predicted KIF5B-KLC1 local fold was used to placed motor domains, CCs, TPRs and mScarlet β-barrel fold (**Fig. S4E-F**; **Fig. S7**). The CC0 dimer, CC2^KHC^-CC3 turn-around and C-terminal tail of KHCs and KLCs were modeled manually. All placed models were real space refined in PHENIX and validated using MolProbity(*70, 73*).

### AlphaFold3 model predictions

To determine kinesin models, sequences for two copies of the KIF5B and KLC1 and two ATP and Mg molecules were entered into a single multi-subunit determination using the AlphaFold3 server (www.alphafoldserver.com)(55). Values of moderate to high confidence pIDDT values per residue displayed (**Fig. S7A**) and their corresponding PAE matrix with accuracy of residue position error. To determine KIF5B-MAP7D3^CT^ AlphaFold3 models, sequences for two copies of KIF5B (residues 1-587) MAP7D3^CT^ (residues 476-876) were entered into a single multi-subunit determination using the AlphaFold3 server. A single representative model is presented in **Fig. S12A** with the moderate to high confidence pIDDT values per residue displayed.

### TIRF assays

TIRF flow chambers were assembled from acid-washed glass coverslips, pre-cleaned slides (ThermoFisher), and double-sided tape. Chambers were functionalized by sequential incubation with 0.5 mg/ml PLL-PEG-biotin (Surface Solutions) for 5–10 min and 0.5 mg/ml streptavidin (ThermoFisher) for 5 min.

Taxol-stabilized MTs (10 µM) in BRB80 buffer (80 mM PIPES pH 6.8, 1 mM MgCl₂, 1 mM EGTA) were flowed into the chamber and incubated for 2–5 min to allow immobilization. Unbound MTs were removed by washing with SRP90 assay buffer (90 mM HEPES, 50 mM KCH₃COO, 2 mM Mg(CH₃COO)₂, 1 mM EGTA, 10% glycerol, pH 7.6) supplemented with 1 mg/ml BSA, 0.05 mg/ml biotin-BSA, 0.2 mg/ml K-casein, 0.5% Pluronic F-127, 10 µM taxol, and 1 µM phalloidin. Finally, Kinesin-1 (KIF5B-KLC1) at the indicated concentrations, with or without MAP7D3^CT^, was introduced in SRP90 assay buffer containing 2 mM Mg-ATP (Sigma).

All imaging was performed on a Nikon TE microscope equipped with a PlanApo 100×/1.49 NA objective, a TIRF illuminator (LU-N4), and an Andor iXon EMCCD camera, controlled with MicroManager 1.4 Software(*74*)). Data were analyzed manually using kymographs generated in ImageJ (Fiji).

## Supplementary Materials

Fig. S1-S12, Legends for movie S1-S3

**Fig. S1:**
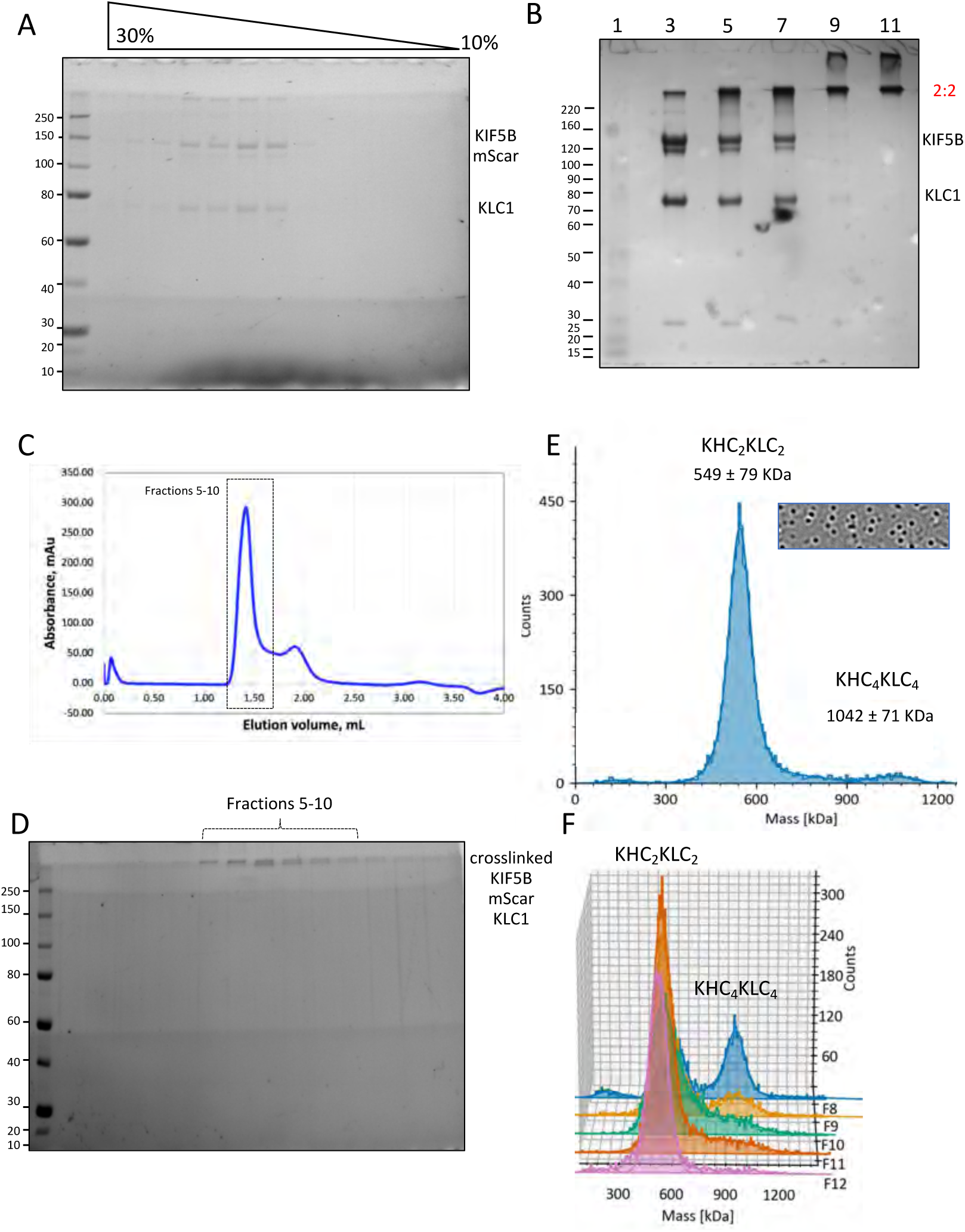
Preparation scheme of Kinesin KIF5B-mScarlet-KLC1 sample for Cryo-EM. A) SDS-PAGE gel showing 10-30% sucrose density gradient of purified KIF5B/KLC1. B) purified KIF5B-KLC1 solution crosslinking experiment with glutaraldehyde. Lane:1 Marker, Lane 3: control KIF5B-mScarlet-KLC1, lane 5: 0.02% glutaraldehyde, lane 7: 0.04% glutaraldehyde, lane 9: 0.08% glutaraldehyde, lane 11: 0.16% glutaraldehyde. Samples were crosslinked at room temperature for 30 minutes and quenched and loaded on the gel to find optimal crosslinker concentration. C) A Size exclusion chromatogram using a Superdex 200 5/150 column to purify glutaraldehyde crosslinked GraFIX KIF5B-mScarlet-KLC1 showing the fractions containing protein (fractions 5-10) corresponding to SDS-PAGE in panel E. D) SDS-PAGE gel showing crosslinked KIF5B/KLC1 sample purified from size exclusion chromatography shown in (D). E) Mass photometry histogram distribution showing mass distribution of GraFIX prepared Kinesin-1 sample shows mostly heterotrimers (KHC_2_KLC_2_) and few larger multimers KHC_4_KLC_4_ particles. F) Mass photometry-based identification of KHC_2_KLC_2_ (heterotetramer) fractions for Cryo-EM analysis by pooling fractions 10-12(F10-F12).

**Fig. S2:**
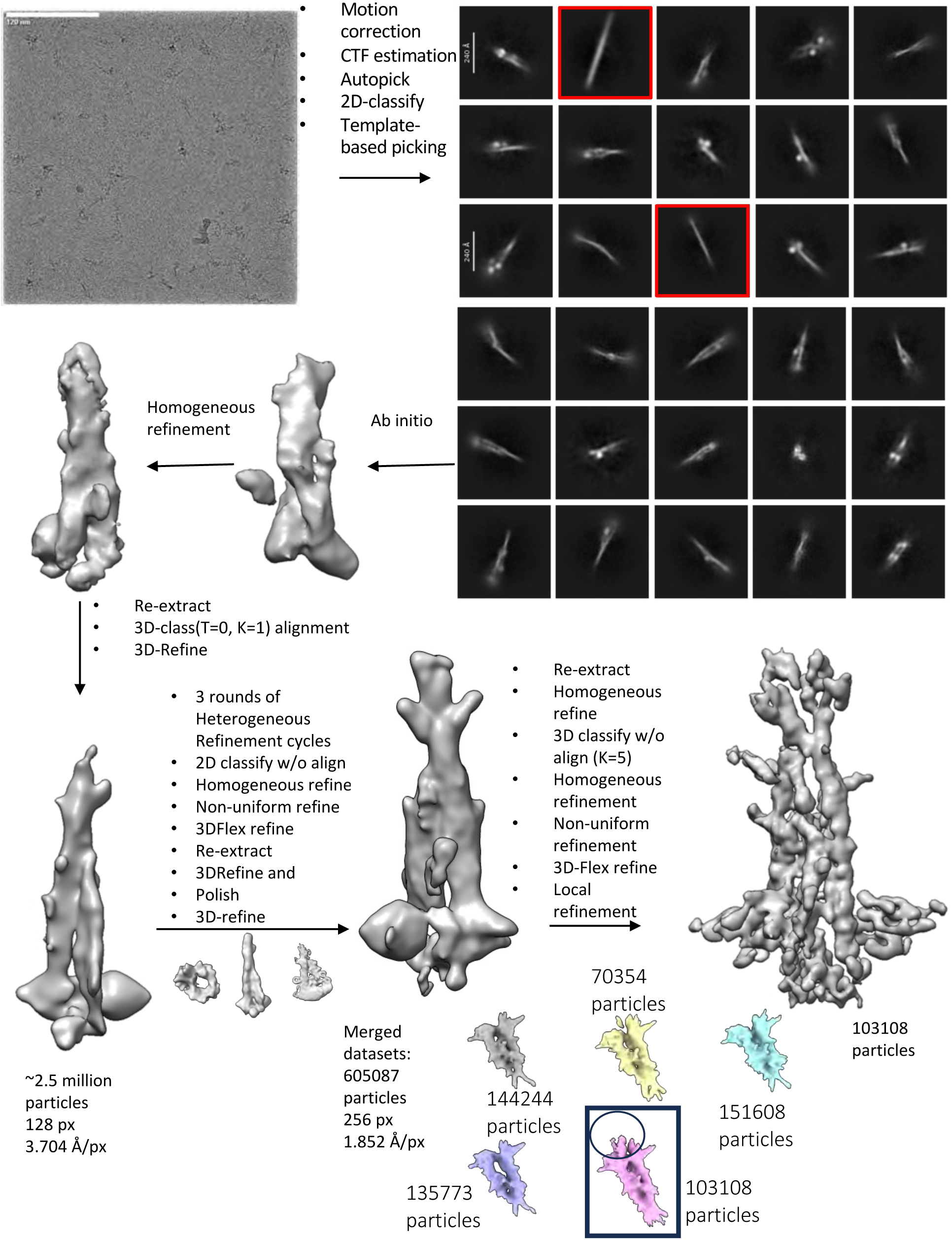
Cryo-EM data processing of KIF5B-mScarlet-KLC1. Cryo-EM movies were collected in EER format which then processed in RELION followed by autopicking and extraction in 200-pixel box (4x binned). 2D classification used for good set of particles and Topaz trained to pick more and 2D classify which then aligned using T=0 classification; Bad classes (highlighted in red) removed Aligned particles were passthrough iterative heterogeneous refine followed by 3D refine and 2D and 3D classification to select final particle set which were then polished and passed through the same cycle as above classification approach.

**Fig. S3:**
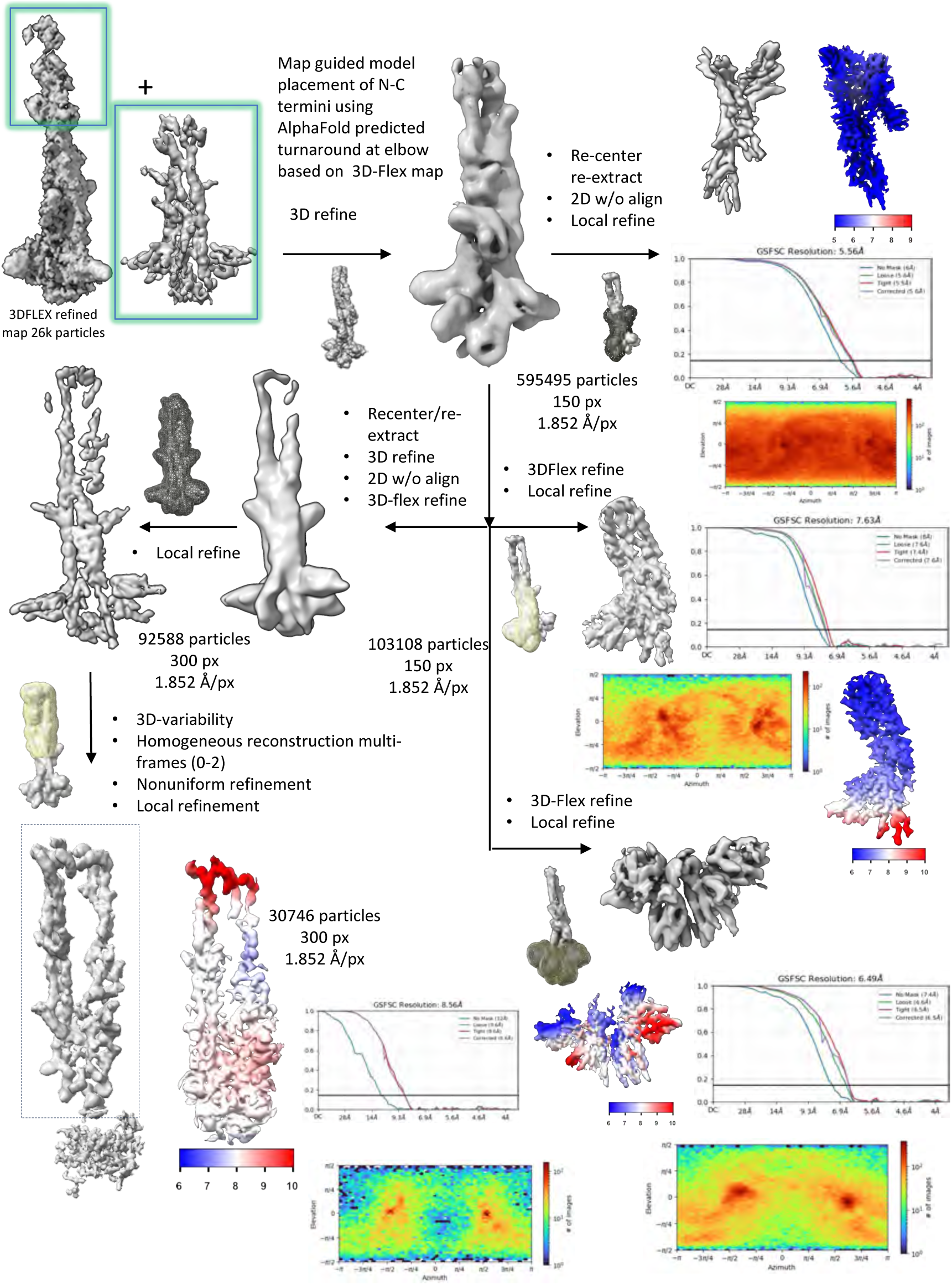
Cryo-EM data processing of KIF5B-mScarlet-KLC1 final refinements. Local refined map and flex refined map were used to place the domains which are then refined against the EMAN generated map lowpass filtered to 15 Å. All particles from four datasets were merged and 3D-refined iteratively followed by 2D without alignment. Placement of model followed by and re-extraction in small box (150 pixel, 2x binned) was local refined for the core of Kinesin-1 consists of KLC (CCs, TPR-B) and KHC (part of CC1^KHC^ and CC4). Merged data was refined and 3D classified without alignment to select 103108 particles with motor density clearly present for both KHCs. Placement of models and re-extraction into small box (150 pix, 2x binned) was used for final local refinements for different parts of motors, TPR(KLC) and tails. Elbow turnaround zone was 3D refined, recenter and reextracted followed by flexible refinement and assignments of densities were used to generate and model-based map to local refine the whole structure was to separate 30k-particles which shows clear turn-around of elbow was 3D-refined by homogeneous reconstruction followed by local refinements.

**Fig. S4:**
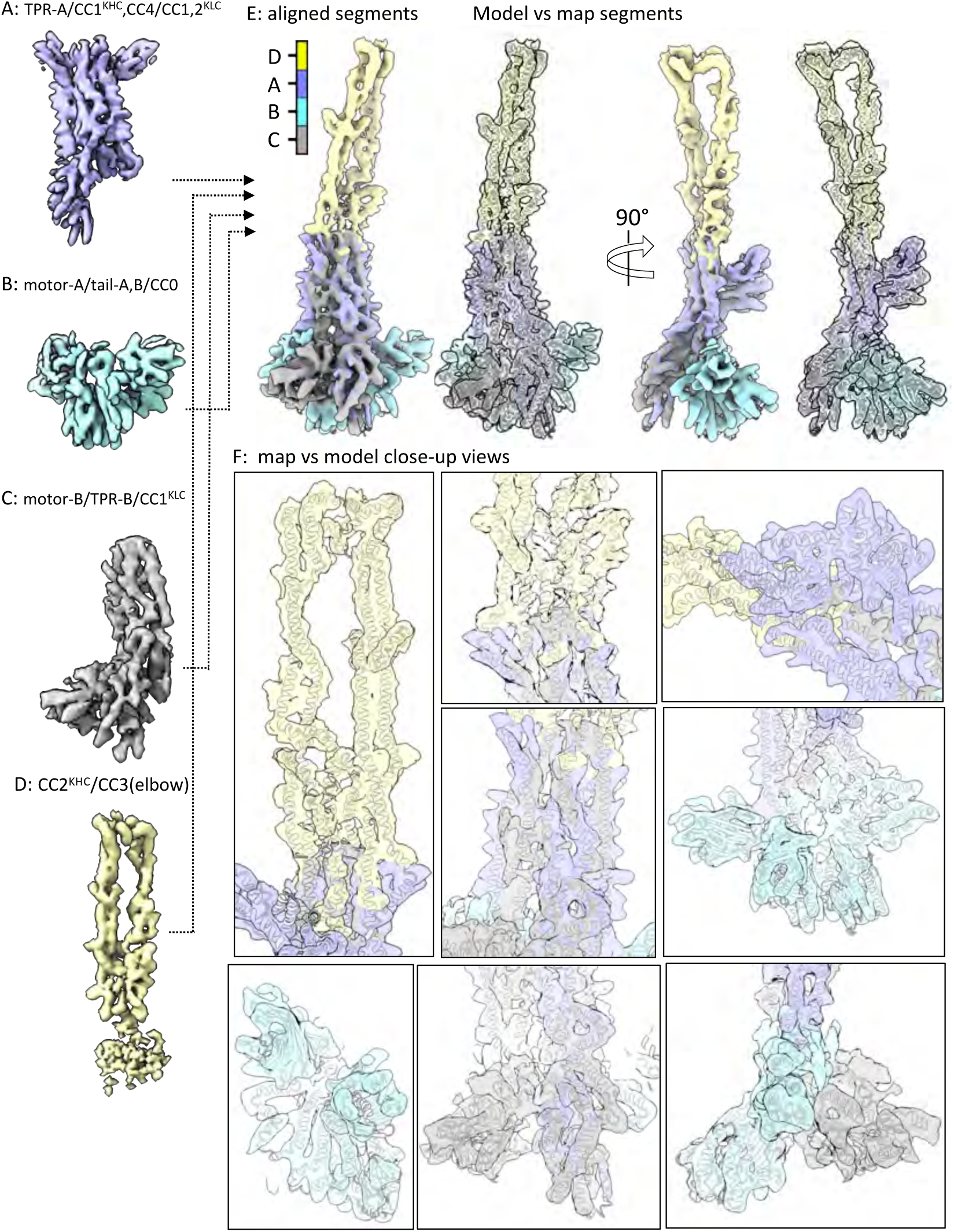
Assembly of regional refined maps to the full kinesin structure map and model. A) Refined subregional map for TPR-A/CC1^KHC^/CC4/CC1-2^KLC^ B) Refined subregional map for motor-A/tail-A, B/CC0 C) Refined subregional map for motor-B/TPR-B/CC1^KHC^ D) Refined subregional map for CC2KHC/CC3 (elbow) E) Left, views of the assembly of all four subregions into the full kinesin map compared to transparent map to model (white ribbon). Right, 90° rotated view of the assembly of all four subregions into the full kinesin map compared to the transparent map to model (white ribbon). A 360° view is also shown in movie S1. F) Close-up views of the map (transparent) to model (white ribbon) for kinesin structure. Top left, helical coiled-coil regions and elbow turn-around zone. Top middle, two overlapping central views CC1^KHC^, CC4, CC1,2^KLC^, Top right, view of TPR-A banded across CC1^KHC,^ CC2^KLC^ and CC4. Middle left, lower CC4, CC1KHC, CC2KLC, L-H A, B; Right middle panel, view of mScar-A,B, tail-A,-B, motor-B and CC0; Lower left, view of CC4, tail-A,B and mScar-A,B; Lower middle, view of motor-A, TPR-B CC1^KHC^, CC4. Lower right, view of motor-A, B and CC0.

**Fig. S5:**
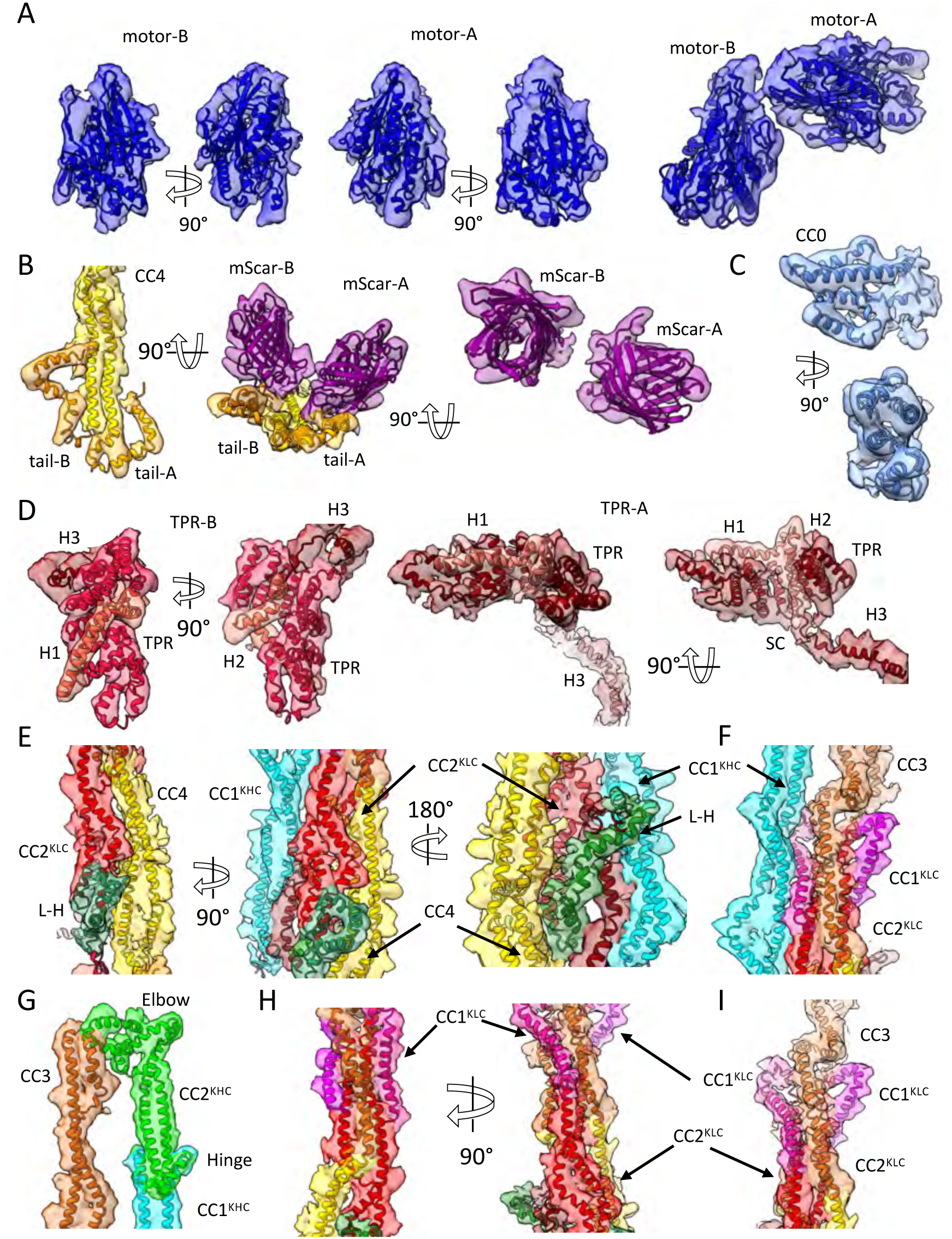
Map and Model close-up showing building of Kinesin-1 domain placements. A) Left, Two rotated views of motor-A domain: Middle, two rotated views of motor-B domain, and a view of both motor-A, B domains. B) Left, end of CC4 and tails map to model showing without mScar-A, B; Middle, 90° rotated CC4 and tail-A, B with mScar-A, B; Right, mScar-A, B β-barrels alone. C) Map to model of CC0 and neck. D) Map to model of TPRs. E) Map to model of central bundle composed of CC1^KHC^ and CC4 and CC1^KLC^, CC2^KLC^ F) Three views of map to model CC1^KHC^-CC3 and CC1-2^KLC^ tetrameric assembly G) Map to model of the elbow structure of CC2^KHC^, CC3. H) Two views of the map to model CC2^KLC^-CC3-CC4 I) side view of the map to model CC1-2^KLC^ CC3-CC4 interface

**Fig. S6:**
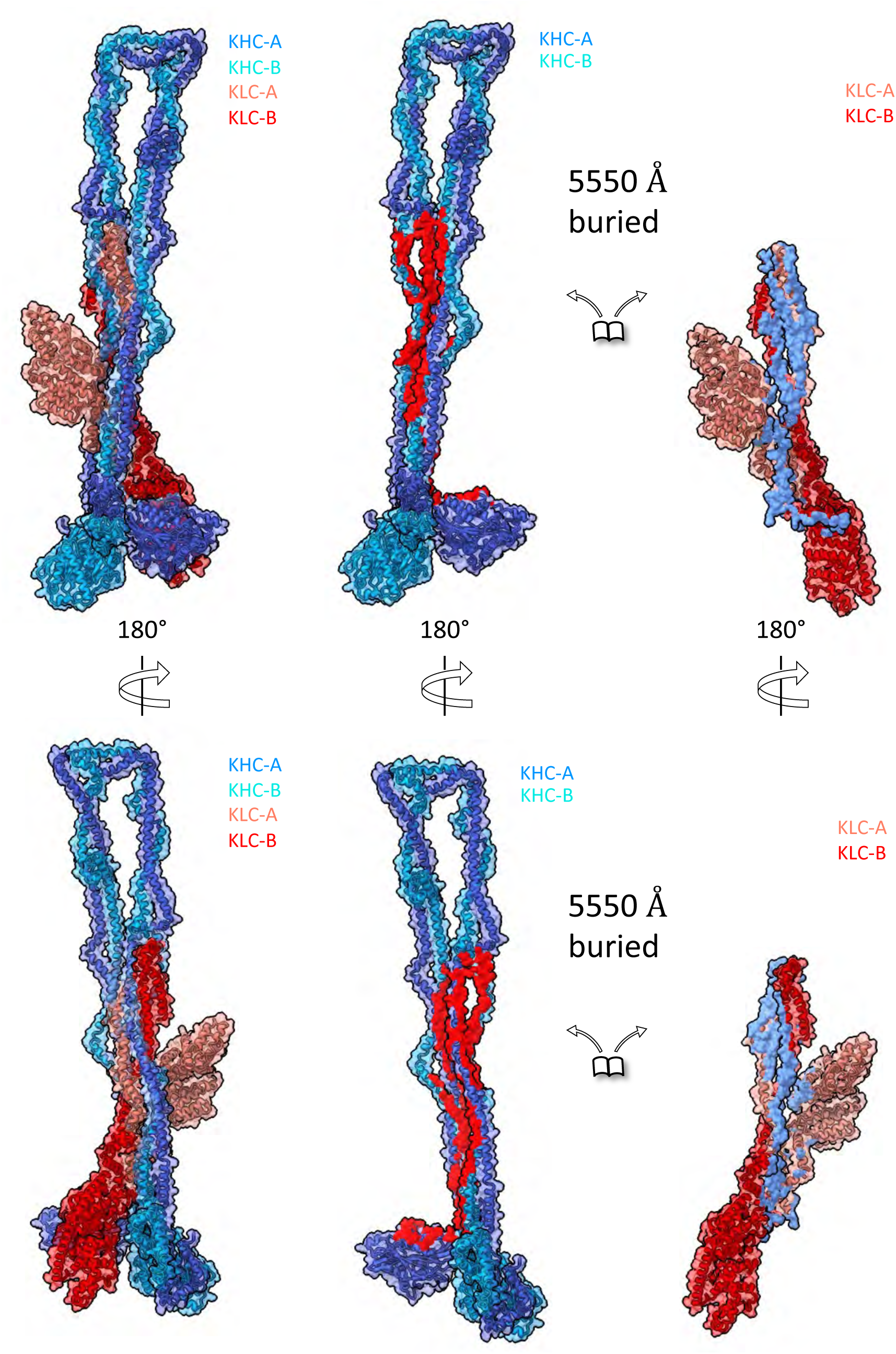
Surface representation showing buried surfaces between KHC and KLC in KIF5B-KLC1 cryo-EM structure. Two views at 180° of KIF5B/KLC1 structure showing the buried surface where at the left full complex is shown and the right KHC and KLC are shown as separate map-to-model in similar color coding. Buried surface area of 5550 Å^2^ is shown in red.

**Fig. S7:**
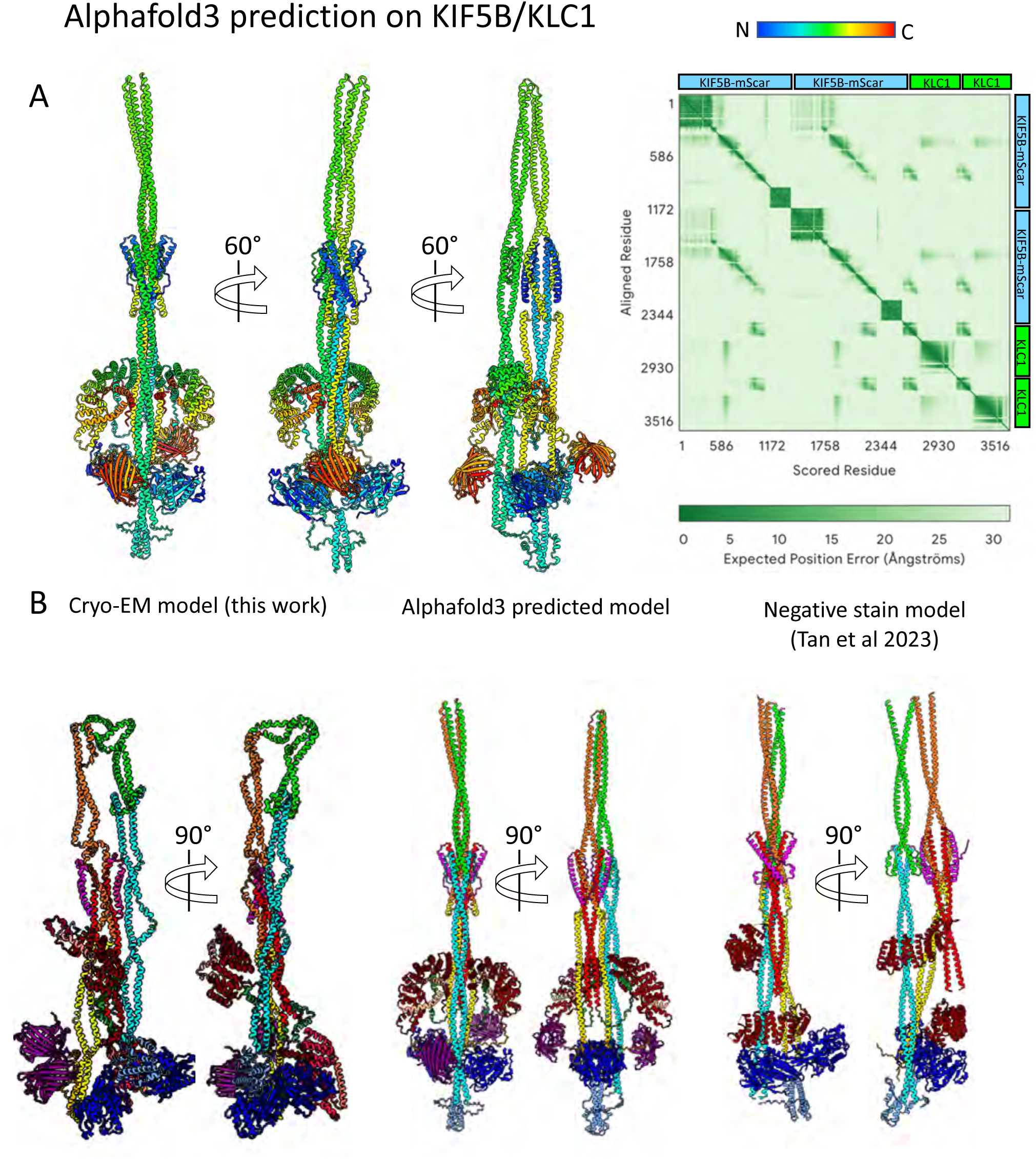
Comparison of Kinesin autoinhibited structures from Cryo-EM, AlphaFold and negative EM structures. A) Left panel, Multiple rotated views of AlphaFold3(*1*) prediction of KIF5B-mScarlet/KLC1 shown in rainbow colors for both KHC and KLC subunits. Right panel, Alphafold3 pLDDT 2D-plot showing the accuracy of domain prediction. B) Top panel, Linear domain representation of KIF5B-KLC1. Bottom panel, Side by side Comparison of Kinesin structural models in consistent color coding as shown in linear domain mapping shown in top pane. Left panels Cryo-EM model based on this study, middle panels the Kinesin-1 Alphafold3 (as shown in A), left panel the negative stain EM model described by Tan et al(*2*). There are no mScarlet proteins fused shown in negative stain model.

**Fig. S8:**
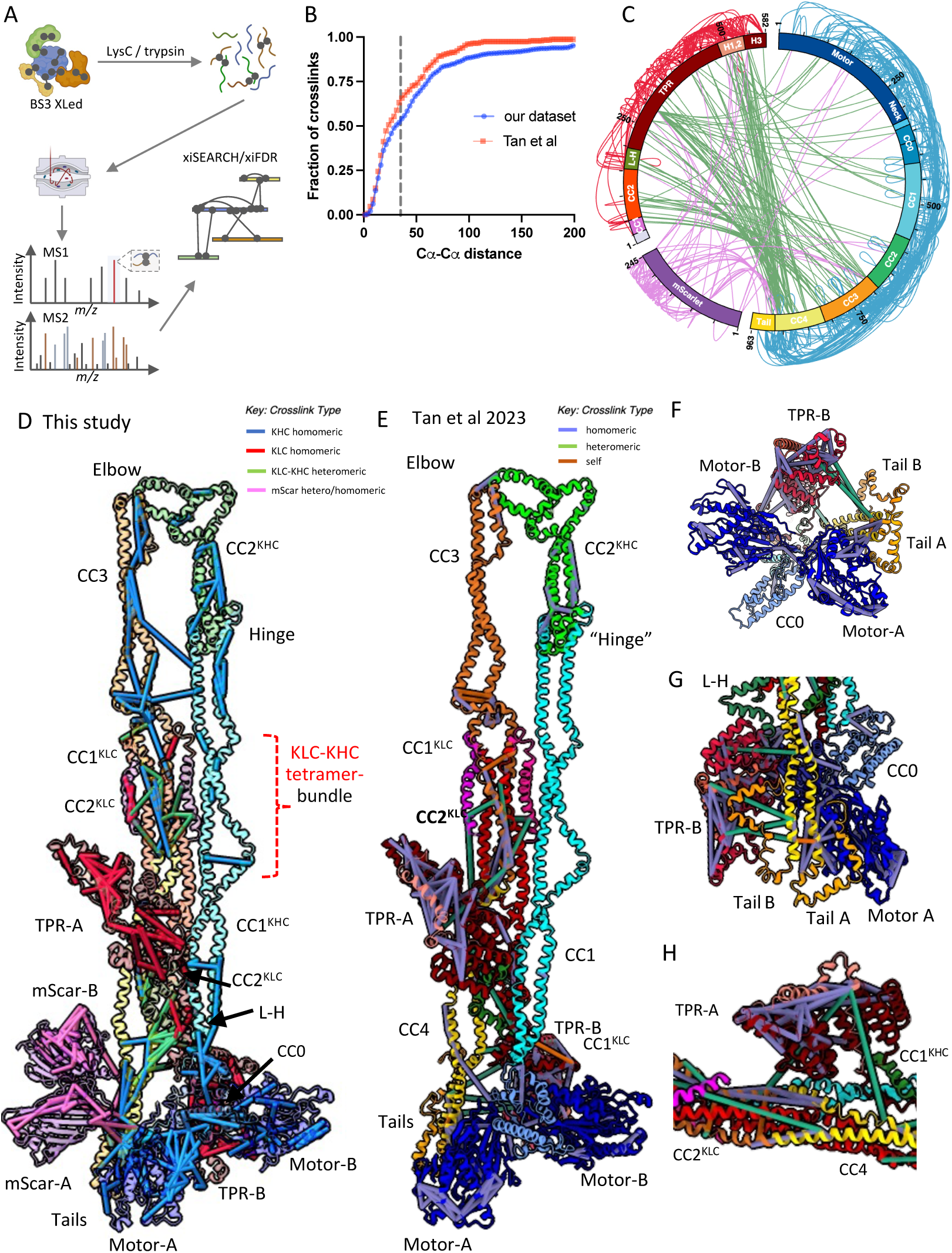
Crosslinking mass spectrometry dataset presented in this study compared to Tan et al. (2) validate the autoinhibited kinesin cryo-EM structure. A) Experimental scheme for crosslinking mass spectrometry. B) CDF plot comparing our dataset with (*2*)(KIF5B-KLC1_pLINK_1), dotted line denotes 35-Å cutoff. C) Connectogram showing all 592 crosslinks identified in our experiment. D) Crosslinks identified in our experiment, illustrated in the context of the cryo-EM structural model of kinesin KIF5B-mScarlet-KLC1 heterotetramer. E) Crosslinks identified in Tan et al (KIF5B-KLC1), illustrated in the context of the cryo-EM structural model of kinesin KIF5B-mScarlet-KLC1 heterotetramer. F) Close-up view of the crosslinks mapped onto motor-A, motor-B, CC0 and KHC tail domains from Tan et al dataset. G) Close up view of the crosslinks mapped onto KHC-CC4, tail-A, tail-B and motor-A and the TPR-B from Tan et al dataset. H) Close up view of the crosslinks mapped onto CC4, CC1^KHC^, CC3 and TPR-A and motor-A and the TPR-B from Tan et al dataset.

**Fig. S9:**
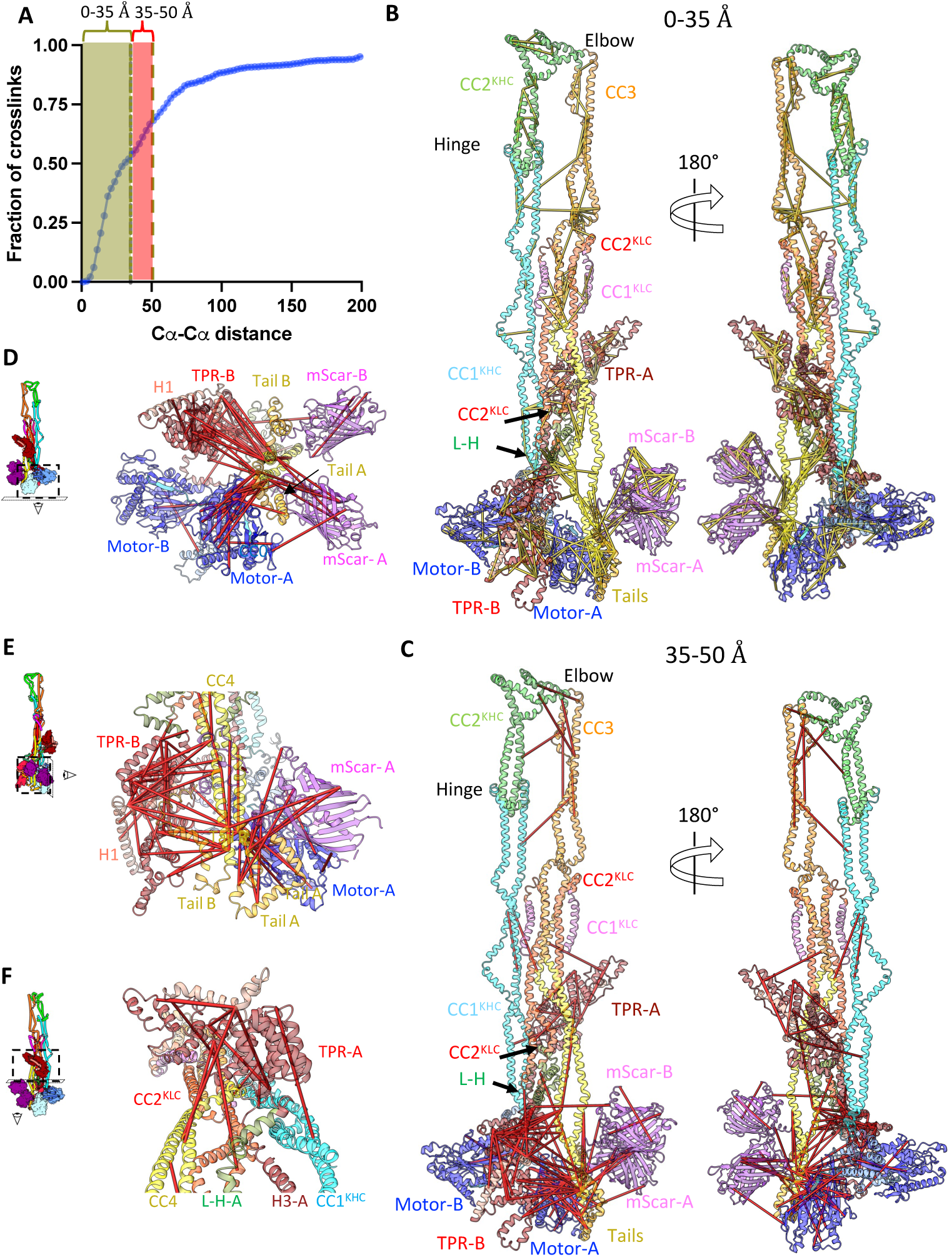
Extended versus normal range distance crosslinks mapped onto the kinesin structure. A) Cumulative distribution function (CDF) plot showing the fraction of the 572 crosslinks whose Cα-Cα distance is 0-35Å distance (yellow region) and crosslinks whose Cα-Cα distance is 36-50Å (red region). The dotted yellow line denotes the 35 Å cutoff and dotted red line denotes the 50Å cutoff. B) Crosslinks (yellow) identified in our experiment, illustrated in the context of the cryo-EM structural model of kinesin KIF5B-mScarlet-KLC1 heterotetramer in two 180° views. In total, 572 crosslinks were successfully mapped onto the kinesin cryo-EM structure, of which 297 with Cα-Cα distances less than 35 Å are displayed. C) Crosslinks (red) identified in our experiment, illustrated in the context of the cryo-EM structural model of kinesin KIF5B-mScarlet-KLC1 heterotetramer in two 180° views. In total, 572 crosslinks were successfully mapped onto the kinesin cryo-EM structure, with Cα-Cα distances 36-50 Å are displayed. D) Bottom view of panel C showing the 36-50Å crosslinks mapped onto KHC motor-A, -B, tail-A, tail-B and the KLC-TPR-B. The view matches that in Fig. 3D. E) Side view of panel C showing the 36-50Å crosslinks mapped onto KHC CC4, tail-A, tail-B and the KLC-TPR-B. The view matches that in Fig. 2C. F) Front view of panel C showing the 36-50Å crosslinks mapped onto KLC-TPR-A, CC2^KLC^, KHC CC1^KHC^, CC4. The view matches that in Fig. 3I.

**Fig. S10:**
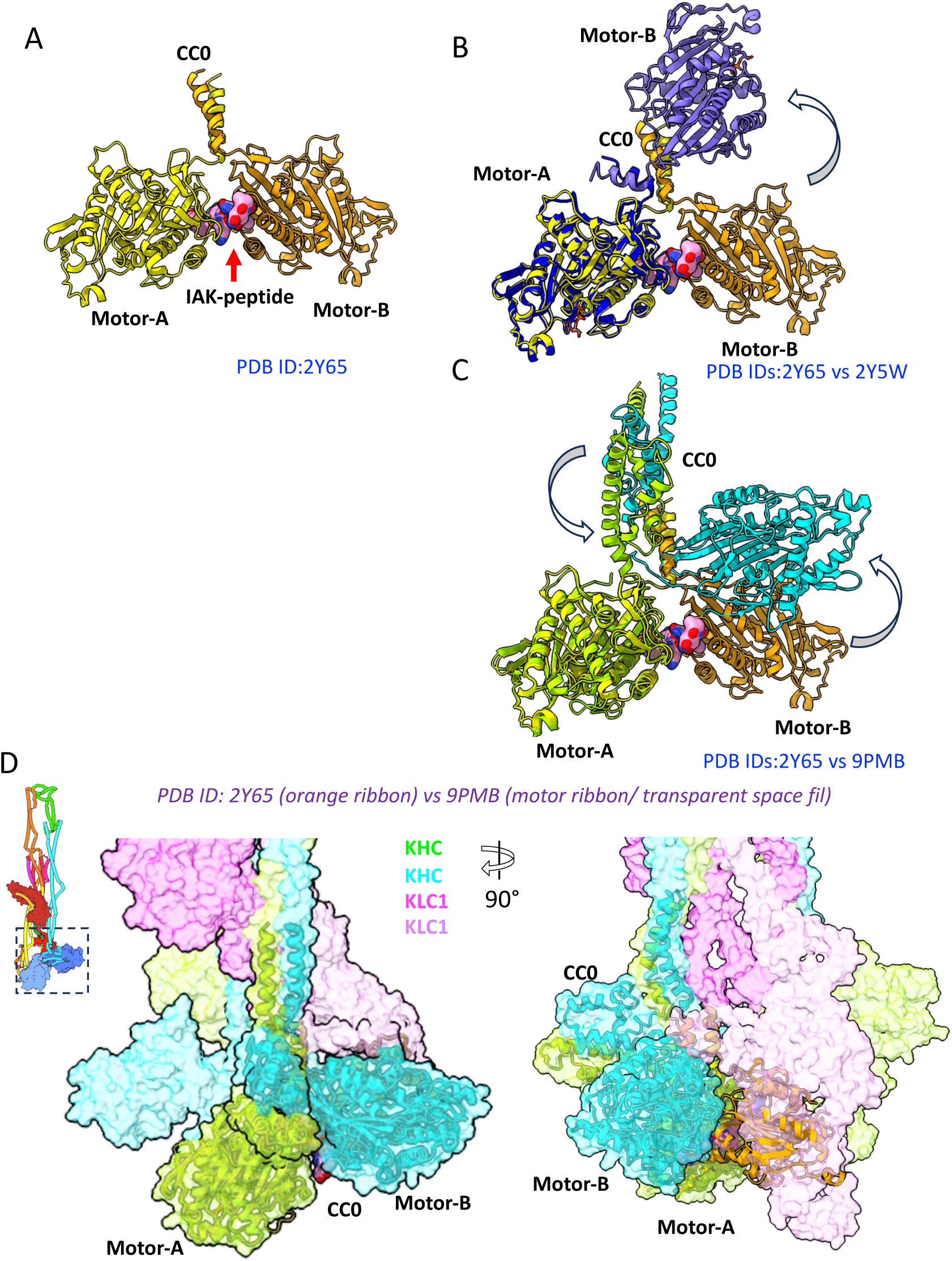
Comparison of motor domains orientations in IAK-peptide binding and cryo-EM model. A) Motor-tail complex (PDB ID:2Y65) is shown where motor domains are colored as yellow (motor-A) and gold (motor-B) and IAK-peptide is shown as pink sphere. B) Comparison of crystal structure of the KHC motor-tail complex (PDB ID: 2Y65) with motor dimer only (PDB ID:2Y5W) as both motor domains (blue). Note motor-B moves outward creating a gap between two motor domains. C) Comparison between the KHC motor-tail interface in autoinhibited kinesin structure presented here compared to KHC motor-tail complex (pdb:2Y65) shown without other domains showing that motor-B is oriented away from motor-A due to TPR-B D) Comparison of cryo-EM model to the motor-tail complex (PDB ID: 2Y65) aligned with motor-A. Note motor-B in crystal structure occupies TPR-B in cryo-EM structure. Cryo-EM model (transparent surface) with KHCs shown as green and KLCs shown as pink and only secondary structures shown for motor-tail assembly. Right panel, a 90° rotated view is shown.

**Fig. S11:**
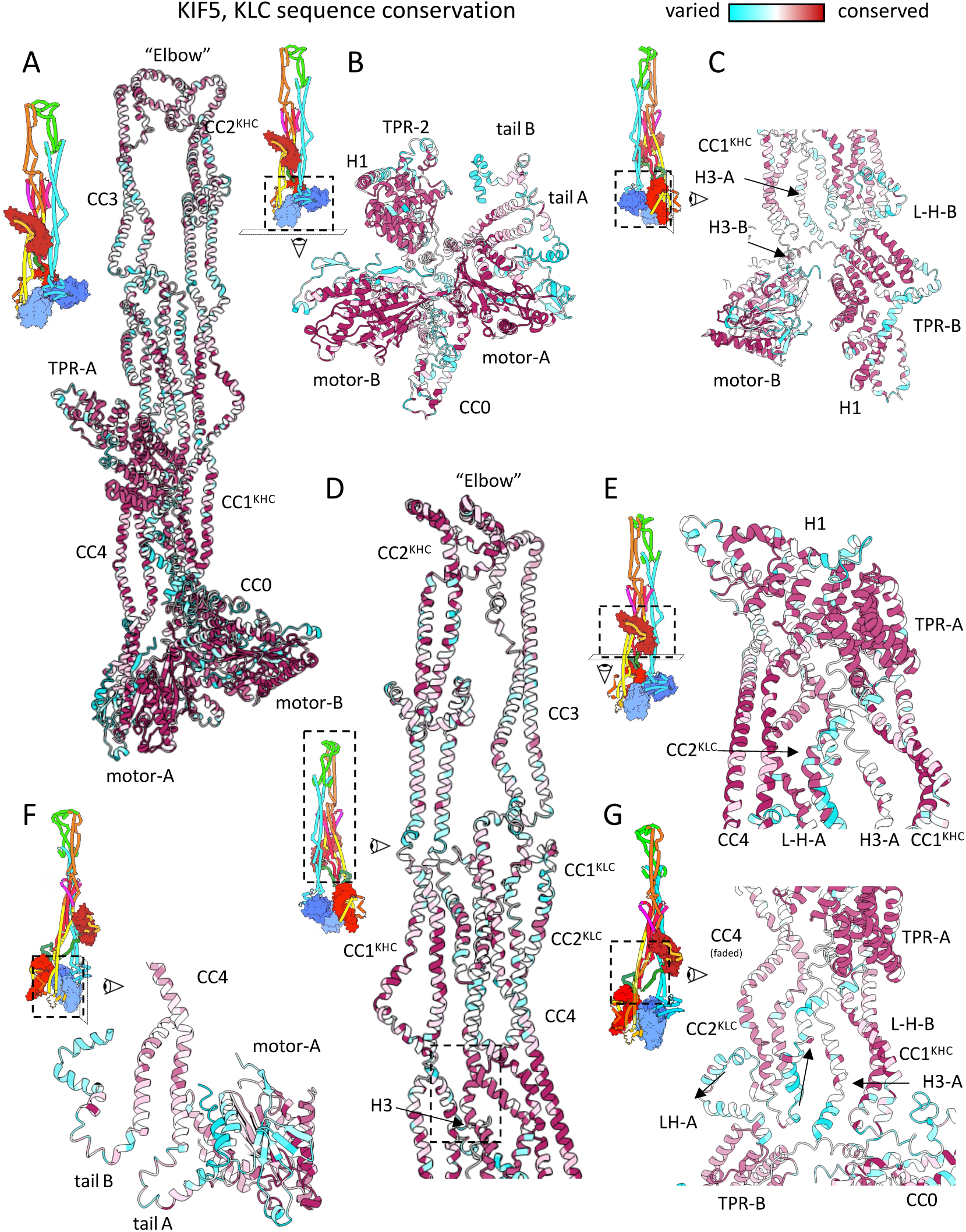
Conservation plot of Kinesin, accompanying close-up views in Fig. 2. A) Full Conservation plot of Kinesin heterotetramer showing the full autoinhibited structure. B) Close-up plot showing motor-tail-TPR interfaces. The motor domains are very conserved with some moderate conservation in CC0 but highly variable C-terminal tails. TPR domains are highly conserved. C) Close up view of Conservation plot showing interfaces of TPR-B/motor-B/H3 where the KLC L-H and H1 are less conserved but H3 is moderately conserved. D) Conservation plot of KHC and KLC coiled-coils showing the interfaces for self-folding are relatively well conserved. The interface of H3/CC1^KHC^/CC1-2^KLC^/CC4 coiled-coils (8 helical bundle) is very conserved. Interface of CC1^KLC^-CC4 is well conserved on the interacting sites. elbow structure is very conserved. E) Conservation plot of KLC-TPR-A on CC4-CC1^KHC^ is in interface with high conservation while the H3 is variable. F) Conservation plot showing KHC motor-A and C-terminal tails showing tails are variable, but their regions that bind the motor domains are well-conserved site while their C-termini are less conserved. G) Conservation plot of the central core showing the KHC coiled-coils and KLC-TPR are well conserved. H3 is partially conserved.

**Fig. S12:**
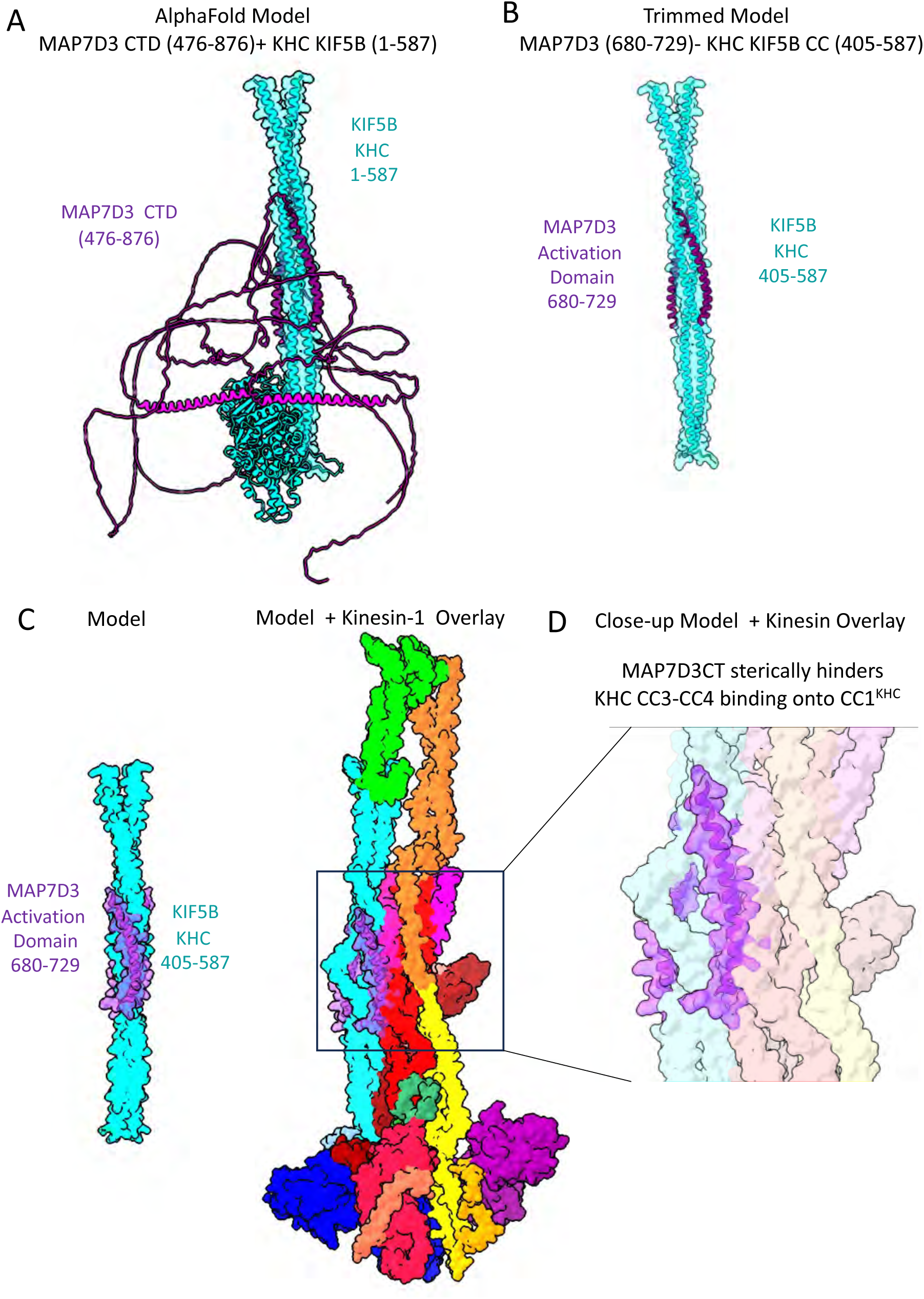
identifying the MAP7D3 binding site on the KHC KIF5B using Alphafold3. A) Alphafold3 results of the MAP7D3CT dimer with KHC KIF5B residues 1-587 B) Trimmed model reveals that MAP7D3 residues 680-729 binds to CC1KHC 405-587. C) Comparison of trimmed model in B to overlaid MAP7D3 model onto Kinesin-1 autoinhibited structure reveals MAP7D3 sterically competes with KHC-CC3-CC4 /CC2KLC binding onto CC1KHC. D) Close up of the boxed view in C, matching Fig. 6A.

## Legends for Movies S1-S3

**Movie S1**: A 360° view of Flexible refine showing kinesin flexibility around CC1^KHC^-CC4 interfacing with KLC and TPR binding to KHC/KLC coiled-coils, followed by a 360°view of the aligned subregional maps with the *de novo* built kinesin heterotetramer ribbon model.

**Movie S2 (accompanies Fig. 1)**: A 360° rotation of the segmented autoinhibited kinesin cryo-EM map with the subunit colors as described in Fig. 1A. This is followed by 360° rotation of the segmented and modeled kinesin cryo-EM map with all subunits colored as in Fig. 1C. This is followed by 360° view of the dissociated view of the modeled kinesin heavy chain (KHC) dimer and kinesin light chain (KLC) dimer colored as shown **Fig. 1D**.

**Movie S3 (accompanies Fig. 2):** A 360° view of the autoinhibited kinesin model with domains colored as presented in **Fig. 2B**.

## Notes

### Competing Interest Statement

The authors have declared no competing interest.

### Summary of Updates

The revised version includes structural validation experimental data and analyses using crosslinking mass spectrometry

