## Supplementary material for "Structural Basis of Kinesin-1 Autoinhibition and Its Control of Microtubule-Based Motility": no links necessary

**Supplementary Materials**

**Fig. S1-S12, Legends for movie S1-S3**

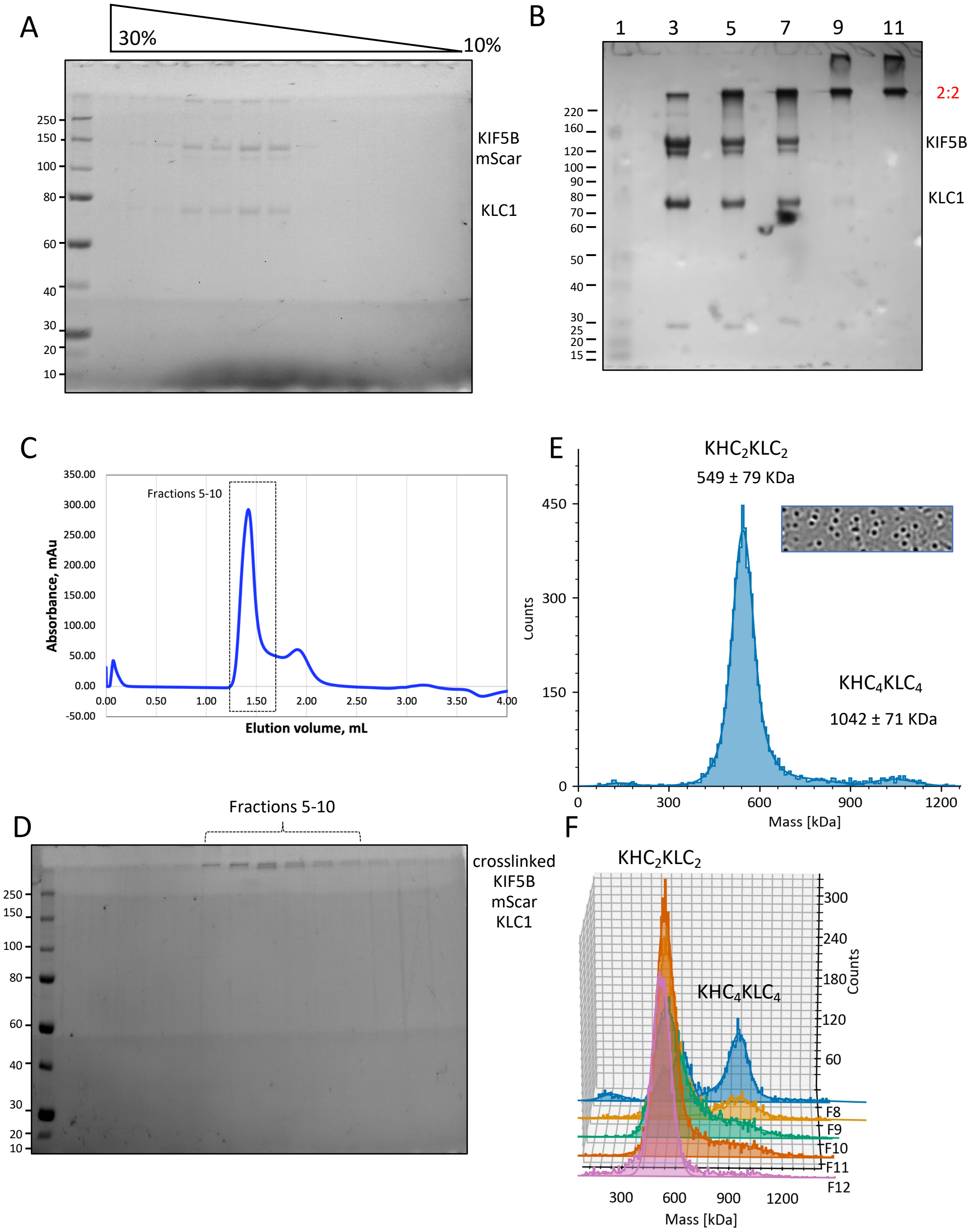


**Fig. S1: Preparation scheme of Kinesin KIF5B-mScarlet-KLC1 sample for Cryo-EM.**

A) SDS-PAGE gel showing 10-30% sucrose density gradient of purified KIF5B/KLC1.

B) purified KIF5B-KLC1 solution crosslinking experiment with glutaraldehyde. Lane:1 Marker, Lane 3: control KIF5B-mScarlet-KLC1, lane 5: 0.02% glutaraldehyde, lane 7: 0.04% glutaraldehyde, lane 9: 0.08% glutaraldehyde, lane 11: 0.16% glutaraldehyde. Samples were crosslinked at room temperature for 30 minutes and quenched and loaded on the gel to find optimal crosslinker concentration.

C) A Size exclusion chromatogram using a Superdex 200 5/150 column to purify glutaraldehyde crosslinked GraFIX KIF5B-mScarlet-KLC1 showing the fractions containing protein (fractions 5-10) corresponding to SDS-PAGE in panel E.

D) SDS-PAGE gel showing crosslinked KIF5B/KLC1 sample purified from size exclusion chromatography shown in (D).

E) Mass photometry histogram distribution showing mass distribution of GraFIX prepared Kinesin-1 sample shows mostly heterotrimers ($KHC_2KLC_2$) and few larger multimers $KHC_4KLC_4$ particles.

F) Mass photometry-based identification of $KHC_2KLC_2$ (heterotetramer) fractions for Cryo-EM analysis by pooling fractions 10-12(F10-F12).

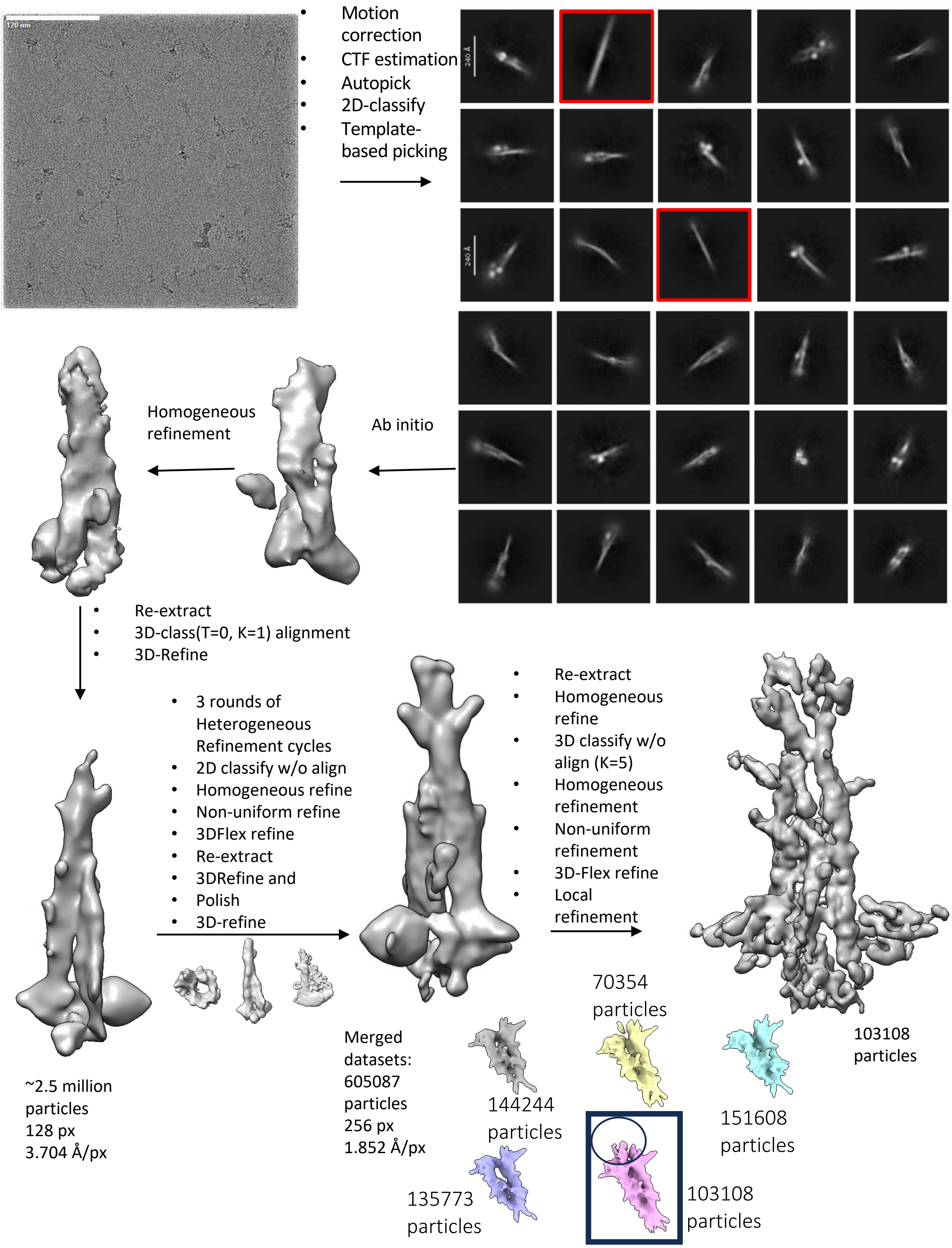


**Fig. S2: Cryo-EM data processing of KIF5B-mScarlet-KLC1.**
Cryo-EM movies were collected in EER format which then processed in RELION followed by autopicking and extraction in 200-pixel box (4x binned). 2D classification used for good set of particles and Topaz trained to pick more and 2D classify which then aligned using T=0 classification; Bad classes (highlighted in red) removed Aligned particles were passthrough iterative heterogeneous refine followed by 3D refine and 2D and 3D classification to select final particle set which were then polished and passed through the same cycle as above classification approach.

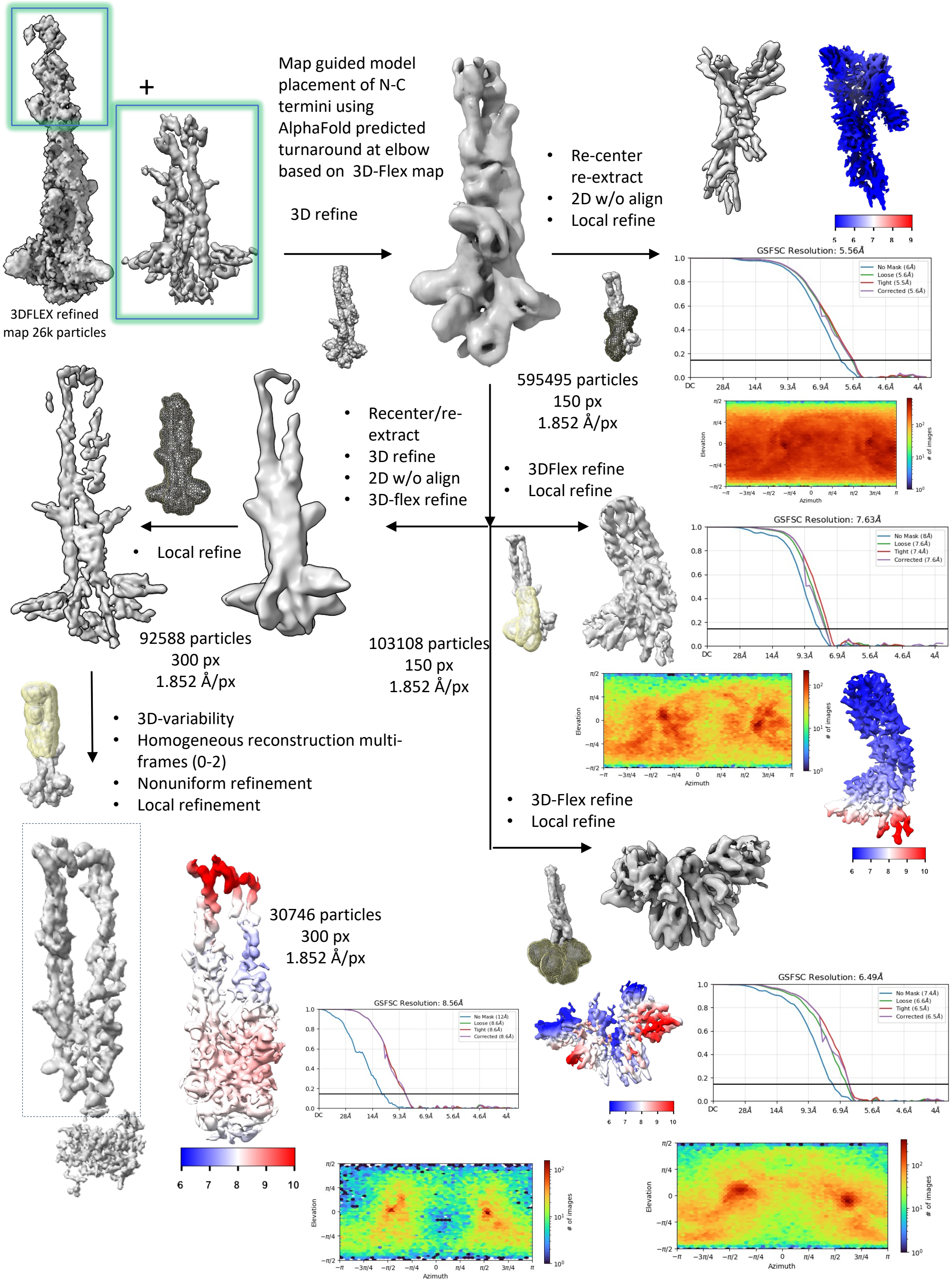


**Fig. S3: Cryo-EM data processing of KIF5B-mScarlet-KLC1 final refinements.**

Local refined map and flex refined map were used to place the domains which are then refined against the EMAN generated map lowpass filtered to 15 Å. All particles from four datasets were merged and 3D-refined iteratively followed by 2D without alignment. Placement of model followed by and re-extraction in small box (150 pixel, 2x binned) was local refined for the core of Kinesin-1 consists of KLC (CCs, TPR-B) and KHC (part of CC1$^{KHC}$ and CC4). Merged data was refined and 3D classified without alignment to select 103108 particles with motor density clearly present for both KHCs. Placement of models and re-extraction into small box (150 pix, 2x binned) was used for final local refinements for different parts of motors, TPR(KLC) and tails. Elbow turnaround zone was 3D refined, recenter and reextracted followed by flexible refinement and assignments of densities were used to generate and model-based map to local refine the whole structure was to separate 30k-particles which shows clear turn-around of elbow was 3D-refined by homogeneous reconstruction followed by local refinements.

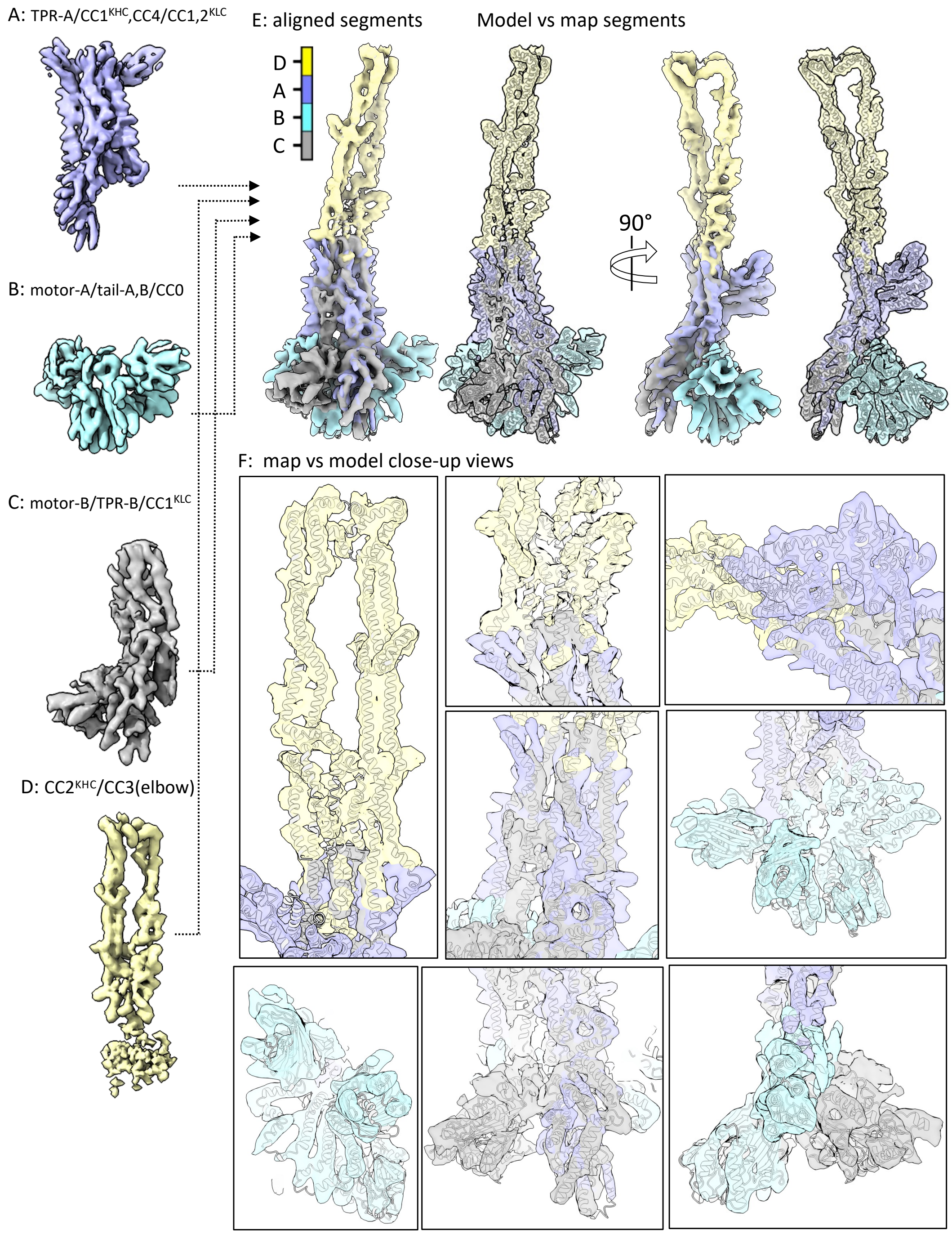


**Fig. S4: Assembly of regional refined maps to the full kinesin structure map and model.**

A) Refined subregional map for TPR-A/CC1$^{KHC}$/CC4/CC1-2$^{KLC}$

B) Refined subregional map for motor-A/tail-A, B/CC0

C) Refined subregional map for motor-B/TPR-B/CC1$^{KHC}$

D) Refined subregional map for CC2KHC/CC3 (elbow)

E) Left, views of the assembly of all four subregions into the full kinesin map compared to transparent map to model (white ribbon). Right, 90° rotated view of the assembly of all four subregions into the full kinesin map compared to the transparent map to model (white ribbon). A 360° view is also shown in movie S1.

F) Close-up views of the map (transparent) to model (white ribbon) for kinesin structure. Top left, helical coiled-coil regions and elbow turn-around zone. Top middle, two overlapping central views CC1$^{KHC}$, CC4, CC1,2$^{KLC}$, Top right, view of TPR-A banded across CC1$^{KHC,}$ CC2$^{KLC}$ and CC4. Middle left, lower CC4, CC1KHC, CC2KLC, L-H A, B; Right middle panel, view of mScar-A,B, tail-A,-B, motor-B and CC0; Lower left, view of CC4, tail-A,B and mScar-A,B; Lower middle, view of motor-A, TPR-B CC1$^{KHC}$, CC4. Lower right, view of motor-A, B and CC0.

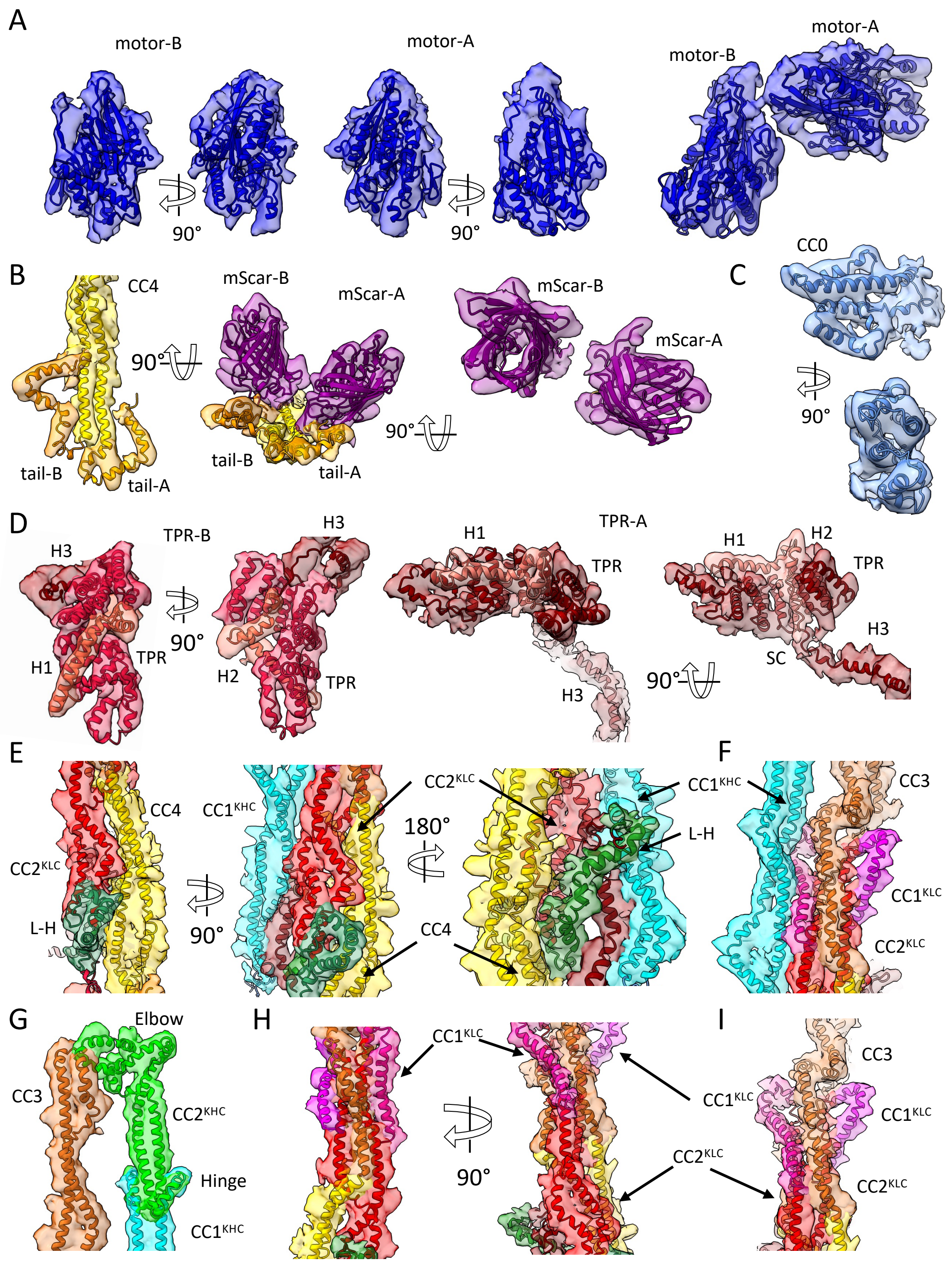


**Fig. S5: Map and Model close-up showing building of Kinesin-1 domain placements.**

A) Left, Two rotated views of motor-A domain: Middle, two rotated views of motor-B domain, and a view of both motor-A, B domains.

B) Left, end of CC4 and tails map to model showing without mScar-A, B; Middle, 90° rotated CC4 and tail-A, B with mScar-A, B; Right, mScar-A, B $\beta$-barrels alone.

C) Map to model of CC0 and neck.

D) Map to model of TPRs.

E) Map to model of central bundle composed of $CC1^{KHC}$ and CC4 and $CC1^{KLC}$, $CC2^{KLC}$

F) Three views of map to model $CC1^{KHC}$-CC3 and $CC1\text{-}2^{KLC}$ tetrameric assembly

G) Map to model of the elbow structure of $CC2^{KHC}$, CC3.

H) Two views of the map to model $CC2^{KLC}$-CC3-CC4

I) side view of the map to model $CC1\text{-}2^{KLC}$ CC3-CC4 interface

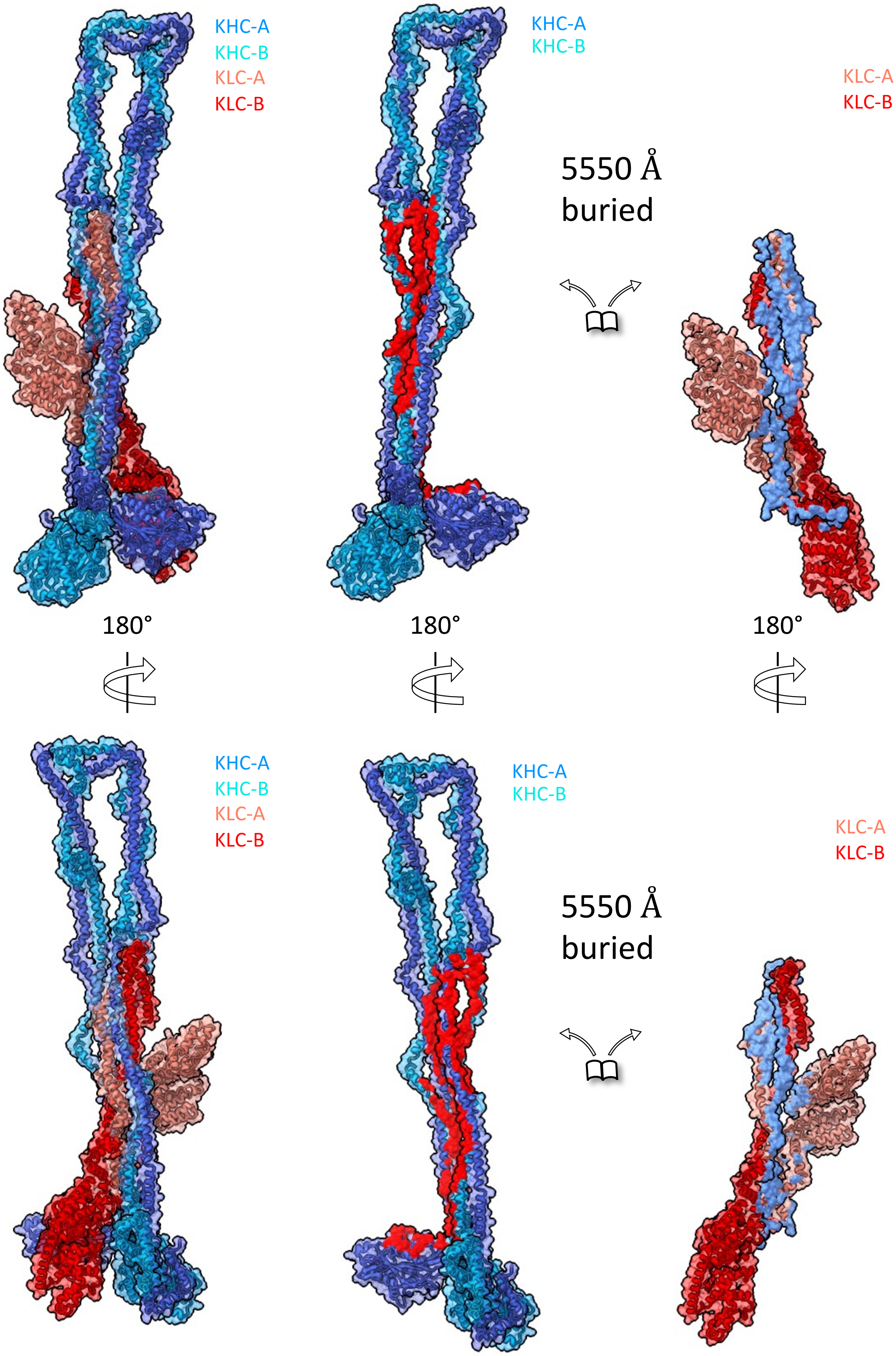


**Fig. S6: Surface representation showing buried surfaces between KHC and KLC in KIF5B-KLC1 cryo-EM structure**.

Two views at 180° of KIF5B/KLC1 structure showing the buried surface where at the left full complex is shown and the right KHC and KLC are shown as separate map-to-model in similar color coding. Buried surface area of 5550 Å$^2$ is shown in red.

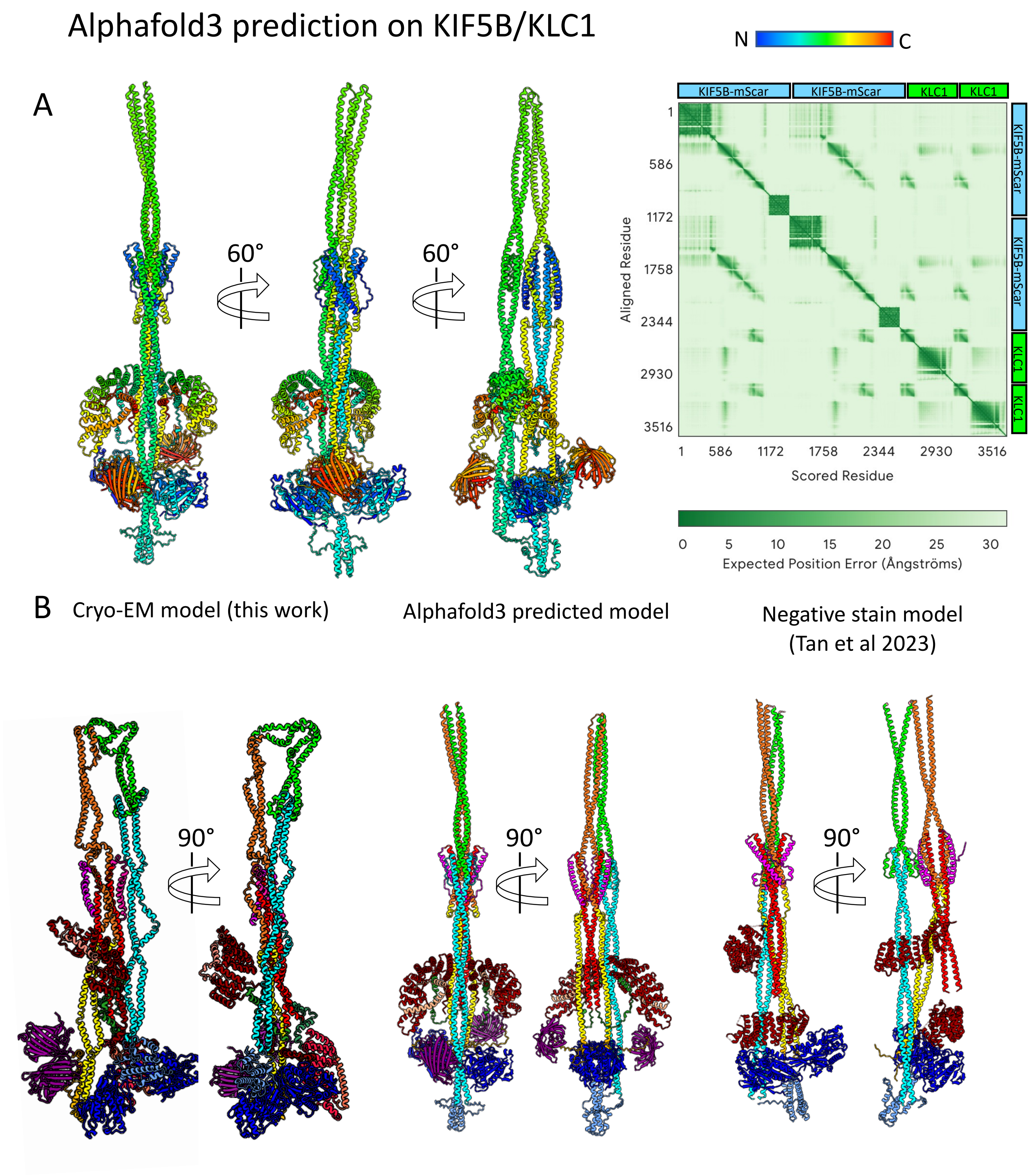


**Fig. S7: Comparison of Kinesin autoinhibited structures from Cryo-EM, AlphaFold and negative EM structures.**

A) Left panel, Multiple rotated views of AlphaFold3(*1*) prediction of KIF5B-mScarlet/KLC1 shown in rainbow colors for both KHC and KLC subunits. Right panel, Alphafold3 pLDDT 2D-plot showing the accuracy of domain prediction.

B) Top panel, Linear domain representation of KIF5B-KLC1. Bottom panel, Side by side Comparison of Kinesin structural models in consistent color coding as shown in linear domain mapping shown in top pane. Left panels Cryo-EM model based on this study , middle panels the Kinesin-1 Alphafold3 (as shown in A), left panel the negative stain EM model described by Tan et al(*2*). There are no mScarlet proteins fused shown in negative stain model.

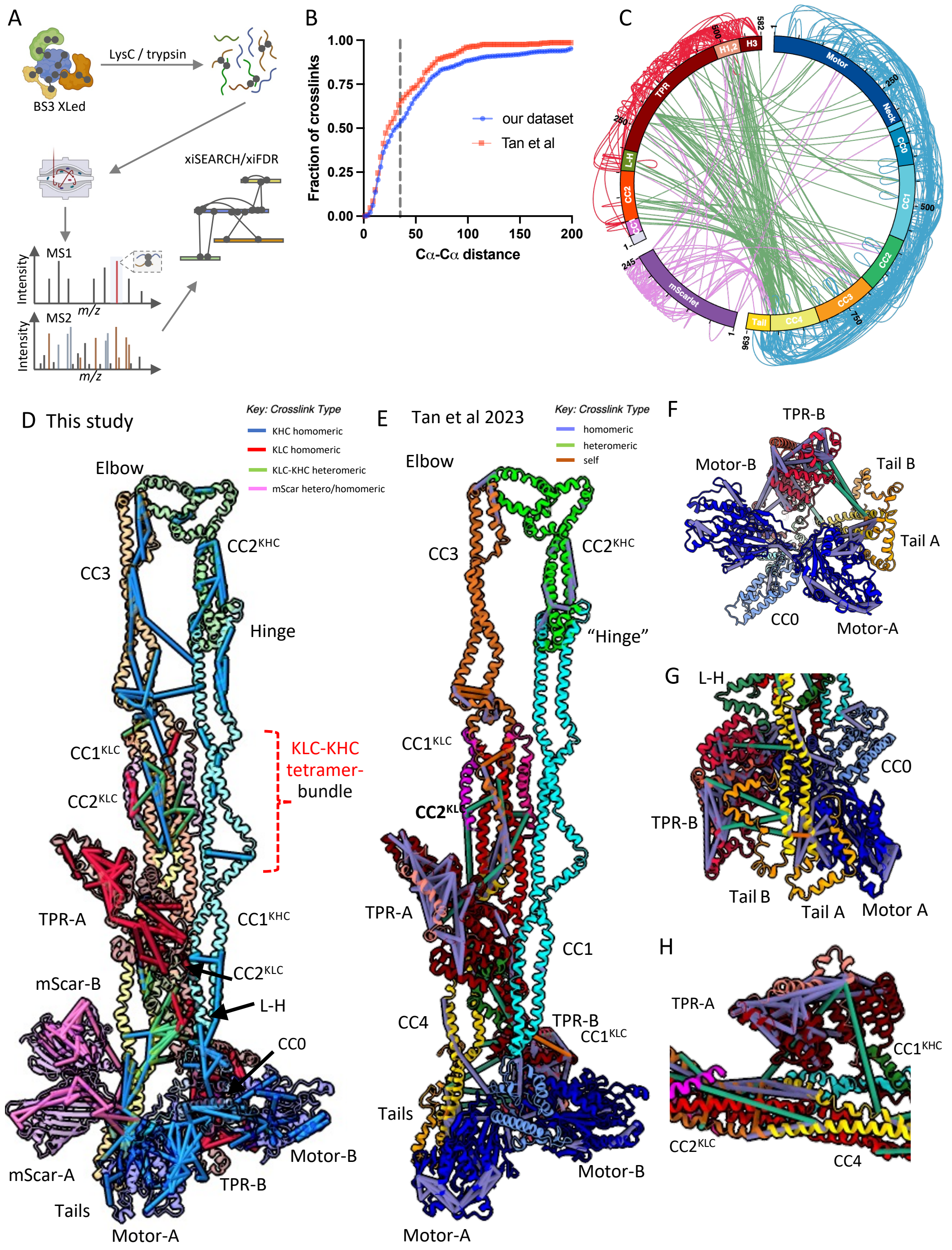


**Fig. S8: Crosslinking mass spectrometry dataset presented in this study compared to Tan et al.(*2*) validate the autoinhibited kinesin cryo-EM structure.**

A) Experimental scheme for crosslinking mass spectrometry.

B) CDF plot comparing our dataset with (*2*)(KIF5B-KLC1_pLINK_1), dotted line denotes 35-Å cutoff.

C) Connectogram showing all 592 crosslinks identified in our experiment.

D) Crosslinks identified in our experiment, illustrated in the context of the cryo-EM structural model of kinesin KIF5B-mScarlet-KLC1 heterotetramer.

E) Crosslinks identified in Tan et al (KIF5B-KLC1), illustrated in the context of the cryo-EM structural model of kinesin KIF5B-mScarlet-KLC1 heterotetramer.

F) Close-up view of the crosslinks mapped onto motor-A, motor-B, CC0 and KHC tail domains from Tan et al dataset.

G) Close up view of the crosslinks mapped onto KHC-CC4, tail-A, tail-B and motor-A and the TPR-B from Tan et al dataset.

H) Close up view of the crosslinks mapped onto CC4, $CC1^{KHC}$, CC3 and TPR-A and motor-A and the TPR-B from Tan et al dataset.

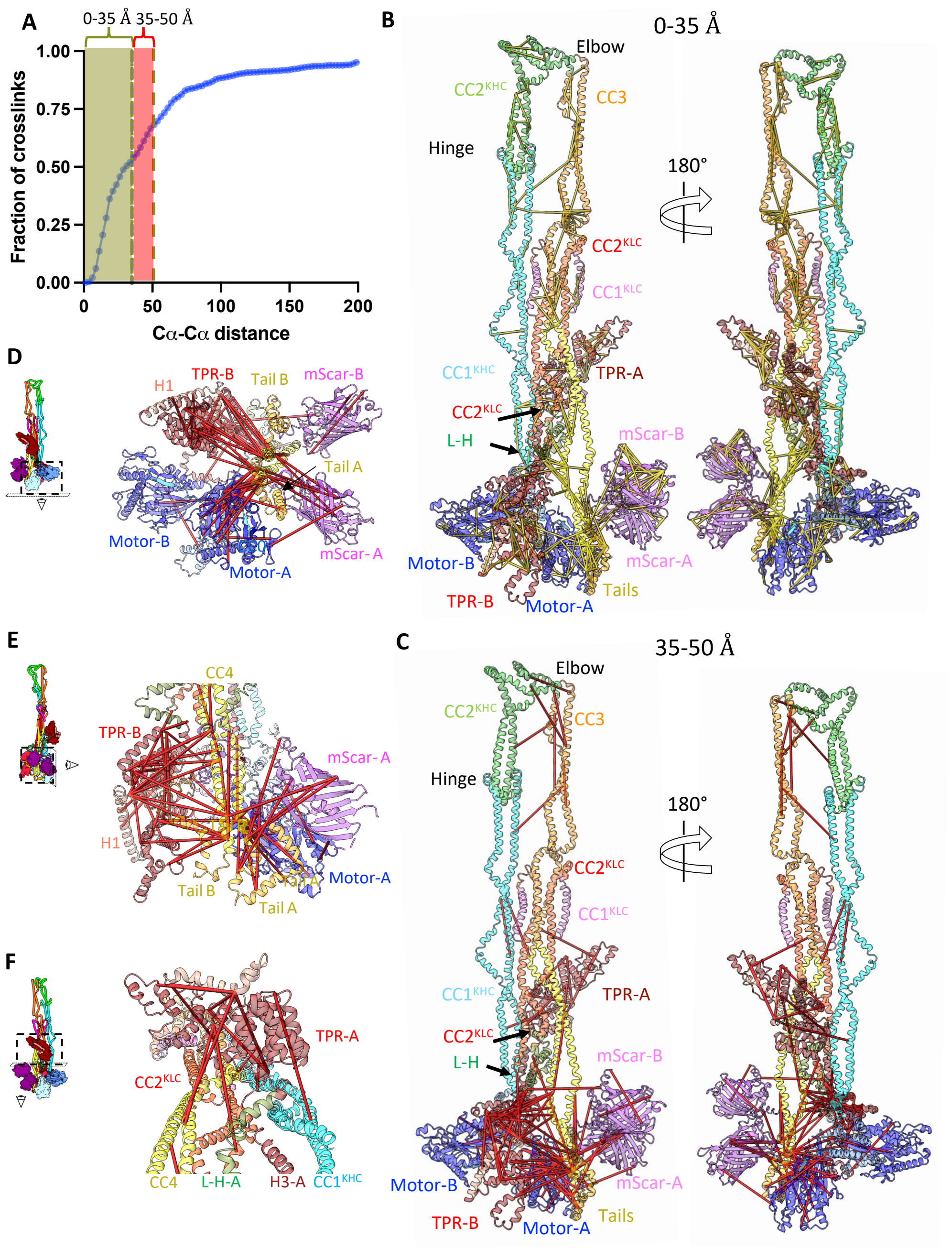


**Fig. S9: Extended versus normal range distance crosslinks mapped onto the kinesin structure**

A) Cumulative distribution function (CDF) plot showing the fraction of the 572 crosslinks whose Cα-Cα distance is 0-35Å distance (yellow region) and crosslinks whose Cα-Cα distance is 36-50Å (red region). The dotted yellow line denotes the 35 Å cutoff and dotted red line denotes the 50Å cutoff.

B) Crosslinks (yellow) identified in our experiment, illustrated in the context of the cryo-EM structural model of kinesin KIF5B-mScarlet-KLC1 heterotetramer in two 180° views. In total, 572 crosslinks were successfully mapped onto the kinesin cryo-EM structure, of which 297 with Cα-Cα distances less than 35 Å are displayed.

C) Crosslinks (red) identified in our experiment, illustrated in the context of the cryo-EM structural model of kinesin KIF5B-mScarlet-KLC1 heterotetramer in two 180° views. In total, 572 crosslinks were successfully mapped onto the kinesin cryo-EM structure, with Cα-Cα distances 36-50 Å are displayed.

D) Bottom view of panel C showing the 36-50Å crosslinks mapped onto KHC motor-A, -B, tail-A, tail-B and the KLC-TPR-B. The view matches that in Fig. 3D.

E) Side view of panel C showing the 36-50Å crosslinks mapped onto KHC CC4, tail-A, tail-B and the KLC-TPR-B. The view matches that in Fig. 2C.

F) Front view of panel C showing the 36-50Å crosslinks mapped onto KLC-TPR-A, $CC2^{KLC}$, KHC $CC1^{KHC}$, CC4. The view matches that in Fig. 3I.

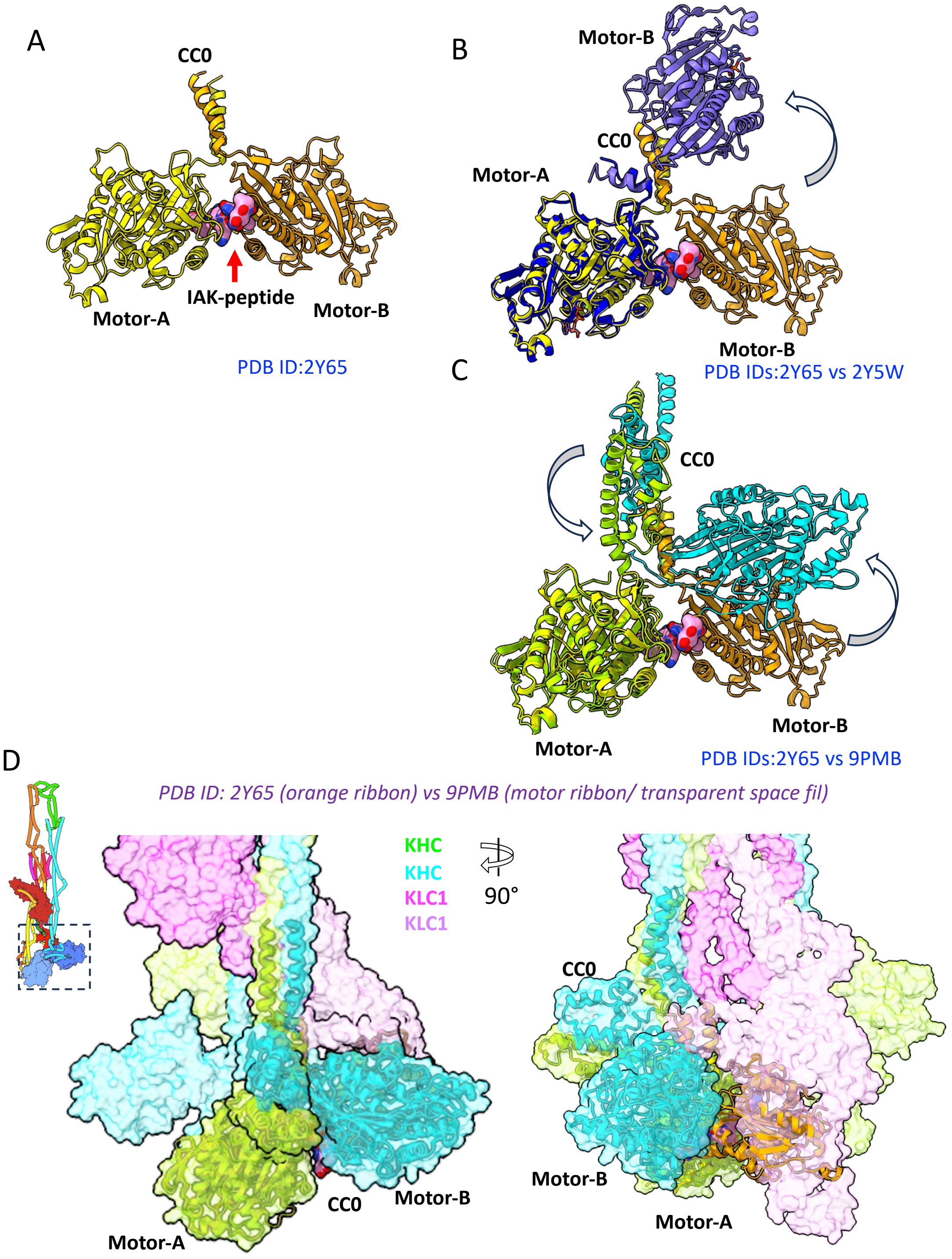


**Fig. S10: Comparison of motor domains orientations in IAK-peptide binding and cryo-EM model.**

A) Motor-tail complex (PDB ID:2Y65) is shown where motor domains are colored as yellow (motor-A) and gold (motor-B) and IAK-peptide is shown as pink sphere.

B) Comparison of crystal structure of the KHC motor-tail complex (PDB ID: 2Y65) with motor dimer only (PDB ID:2Y5W) as both motor domains (blue). Note motor-B moves outward creating a gap between two motor domains.

C) Comparison between the KHC motor-tail interface in autoinhibited kinesin structure presented here compared to KHC motor-tail complex (pdb:2Y65) shown without other domains showing that motor-B is oriented away from motor-A due to TPR-B

D) Comparison of cryo-EM model to the motor-tail complex (PDB ID: 2Y65) aligned with motor-A. Note motor-B in crystal structure occupies TPR-B in cryo-EM structure. Cryo-EM model (transparent surface) with KHCs shown as green and KLCs shown as pink and only secondary structures shown for motor-tail assembly. Right panel, a 90° rotated view is shown.

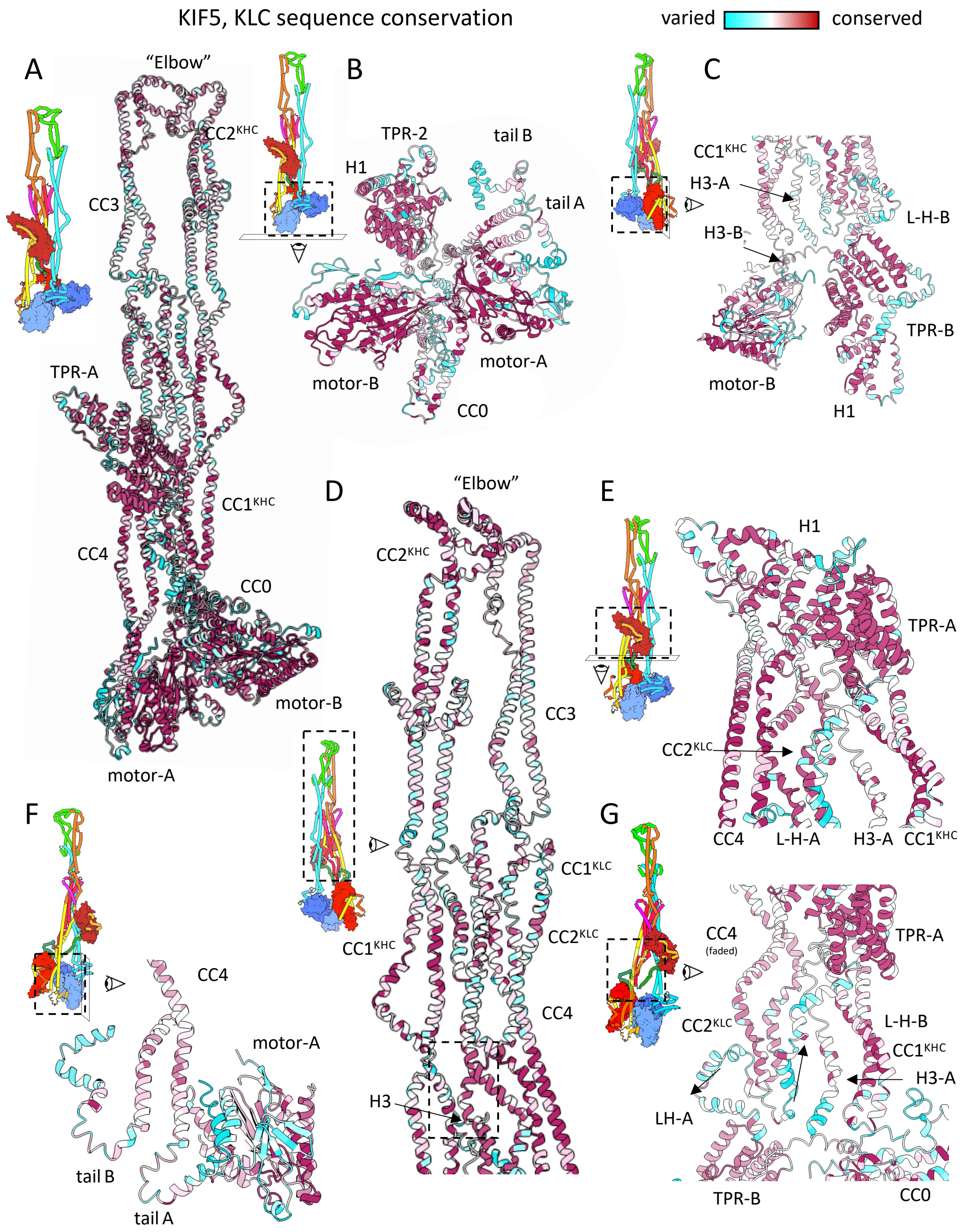


**Fig. S11: Conservation plot of Kinesin, accompanying close-up views in Fig. 2.**

A) Full Conservation plot of Kinesin heterotetramer showing the full autoinhibited structure.

B) Close-up plot showing motor-tail-TPR interfaces. The motor domains are very conserved with some moderate conservation in CC0 but highly variable C-terminal tails. TPR domains are highly conserved.

C) Close up view of Conservation plot showing interfaces of TPR-B/motor-B/H3 where the KLC L-H and H1 are less conserved but H3 is moderately conserved.

D) Conservation plot of KHC and KLC coiled-coils showing the interfaces for self-folding are relatively well conserved. The interface of H3/CC1$^{KHC}$/CC1-2$^{KLC}$/CC4 coiled-coils (8 helical bundle) is very conserved. Interface of CC1$^{KLC}$-CC4 is well conserved on the interacting sites. elbow structure is very conserved.

E) Conservation plot of KLC-TPR-A on CC4-CC1$^{KHC}$ is in interface with high conservation while the H3 is variable.

F) Conservation plot showing KHC motor-A and C-terminal tails showing tails are variable, but their regions that bind the motor domains are well-conserved site while their C-termini are less conserved.

G) Conservation plot of the central core showing the KHC coiled-coils and KLC-TPR are well conserved. H3 is partially conserved.

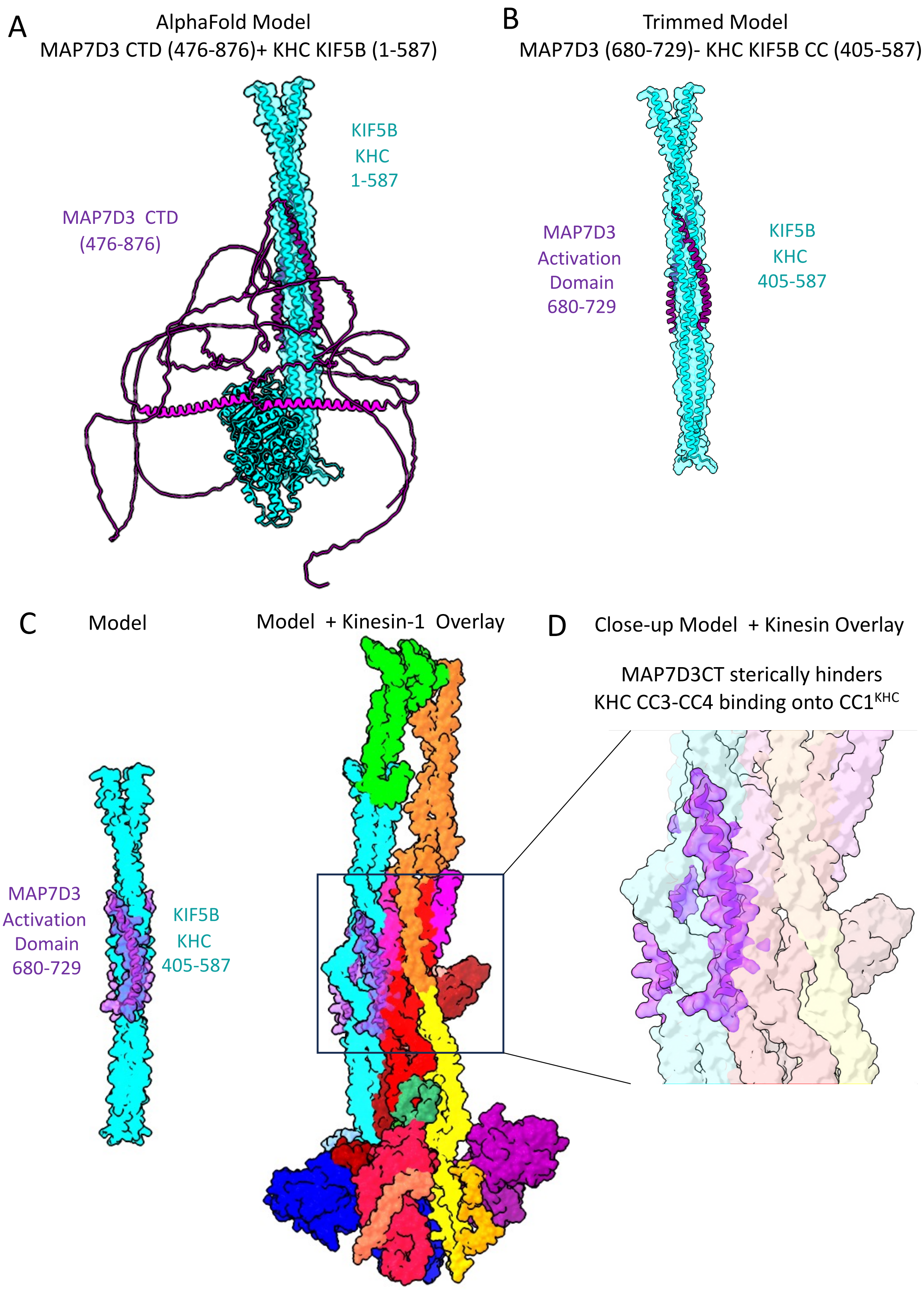


**Fig. S12: identifying the MAP7D3 binding site on the KHC KIF5B using Alphafold3.**

A) Alphafold3 results of the MAP7D3CT dimer with KHC KIF5B residues 1-587

B) Trimmed model reveals that MAP7D3 residues 680-729 binds to CC1KHC 405-587.

C) Comparison of trimmed model in B to overlaid MAP7D3 model onto Kinesin-1 autoinhibited structure reveals MAP7D3 sterically competes with KHC-CC3-CC4 /CC2KLC binding onto CC1KHC.

D) Close up of the boxed view in C, matching Fig. 6A.

**Legends for Movies S1-S3**

**Movie S1**: A 360° view of Flexible refine showing kinesin flexibility around $CC1^{KHC}$-CC4 interfacing with KLC and TPR binding to KHC/KLC coiled-coils, followed by a 360°view of the aligned subregional maps with the *de novo* built kinesin heterotetramer ribbon model.

**Movie S2 (accompanies Fig. 1)**: A 360° rotation of the segmented autoinhibited kinesin cryo-EM map with the subunit colors as described in Fig. 1A. This is followed by 360° rotation of the segmented and modeled kinesin cryo-EM map with all subunits colored as in Fig. 1C. This is followed by 360° view of the dissociated view of the modeled kinesin heavy chain (KHC) dimer and kinesin light chain (KLC) dimer colored as shown **Fig. 1D.**

**Movie S3 (accompanies Fig. 2):** A 360° view of the autoinhibited kinesin model with domains colored as presented in **Fig. 2B**.
